# *Ex vivo* mass cytometry analysis reveals a profound myeloid proinflammatory signature in psoriatic arthritis synovial fluid

**DOI:** 10.1101/2021.03.04.433903

**Authors:** Nicole Yager, Suzanne Cole, Alicia Lledo Lara, Ash Maroof, Frank Penkava, Julian Knight, Paul Bowness, Hussein Al-Mossawi

## Abstract

**Objectives:** A number of immune populations have been implicated in psoriatic arthritis (PsA) pathogenesis. This study used mass cytometry (CyTOF) combined with transcriptomic analysis to generate a high-dimensional dataset of matched PsA synovial fluid (SF) and blood leucocytes, with the aim of identifying cytokine production *ex vivo* in unstimulated lymphoid and myeloid cells.

**Methods:** Fresh SF and paired blood were either fixed or incubated with protein transport inhibitors for 6 h. Samples were stained with two CyTOF panels: a phenotyping panel and an intracellular panel, including antibodies to both T cell and myeloid cell secreted proteins. Transcriptomic analysis by gene array of key expanded cell populations and single-cell RNA-seq, and ELISA and LEGENDplex analysis of PsA SF were also performed.

**Results:** We observed marked changes in the myeloid compartment of PsA SF relative to blood, with expansion of intermediate monocytes, macrophages and dendritic cell populations. Classical monocytes, intermediate monocytes and macrophages spontaneously produced significant levels of the proinflammatory mediators osteopontin and CCL2 in the absence of any *in vitro* stimulation. By contrast minimal spontaneous cytokine production by T cells was detected. Gene expression analysis showed the genes for osteopontin and CCL2 to be amongst those most highly upregulated by PsA monocytes/macrophages; and both proteins were elevated in PsA SF.

**Conclusions:** Using multiomic analyses we have generated a comprehensive cellular map of PsA SF and blood to reveal key expanded myeloid proinflammatory modules in PsA of potential pathogenic and therapeutic importance.

## Introduction

Psoriatic arthritis (PsA) is an immune-mediated inflammatory arthritis which forms part of the spondyloarthropathy (SpA) spectrum. Histopathological characteristics of PsA include enthesitis, synovitis, erosions and new bone formation. The pathogenesis of joint inflammation in PsA is poorly understood;^1^ roles for TNF, IL-17A and IL-23 have been demonstrated with clinical efficacy of neutralising therapies against these cytokine targets.^2-4^ Previous studies, which have predominantly focused on specific immune cell types within PsA synovial fluid (SF), have established potential roles for CD8 T cells,^5^ particularly those producing IL-17,^6^ natural killer (NK) cells^7^ and dendritic cells (DCs).^8^ Other myeloid populations have been relatively under-studied in PsA but research points to the importance of this compartment in PsA pathogenesis. Within the synovium, CD163+ macrophages are increased in SpA compared to RA, despite similar total CD68+ macrophage numbers; and blood-derived CD163+ cells have been shown to produce TNF (following *in vitro* LPS stimulation).^9^

Until recently, comprehensive characterisation of the cellular composition in the psoriatic joint has been technically difficult to achieve. In this study we use mass cytometry (CyTOF) to simultaneously measure over 30 parameters in PsA SF and blood leucocytes directly *ex vivo* and then visualise the inflammatory cellular architecture of PsA using unsupervised clustering. We identify inflammatory proteins including osteopontin and CCL2 spontaneously produced by PsA SF myeloid cells without any *in vitro* stimulation. We combine this with both bulk and single-cell transcriptomic analyses and SF protein quantification to identify a prominent role for myeloid-derived mediators in the pathogenesis of PsA.

## Results

### PsA synovial fluid (SF) shows marked increases in specific myeloid populations compared to PsA blood

CyTOF analysis of matched SF and blood from 11 PsA patients was performed, together with 15 blood samples from healthy donors (workflow outlined in figure 1A). We used a phenotyping panel containing 36 markers to enable identification of all major immune cell populations together with their activation status when samples were fixed immediately (T0). Principal component analysis (PCA) clearly distinguished PsA SF samples from blood; while blood samples from PsA patients and healthy donors were interspersed (figure 1B).

**Figure 1.**
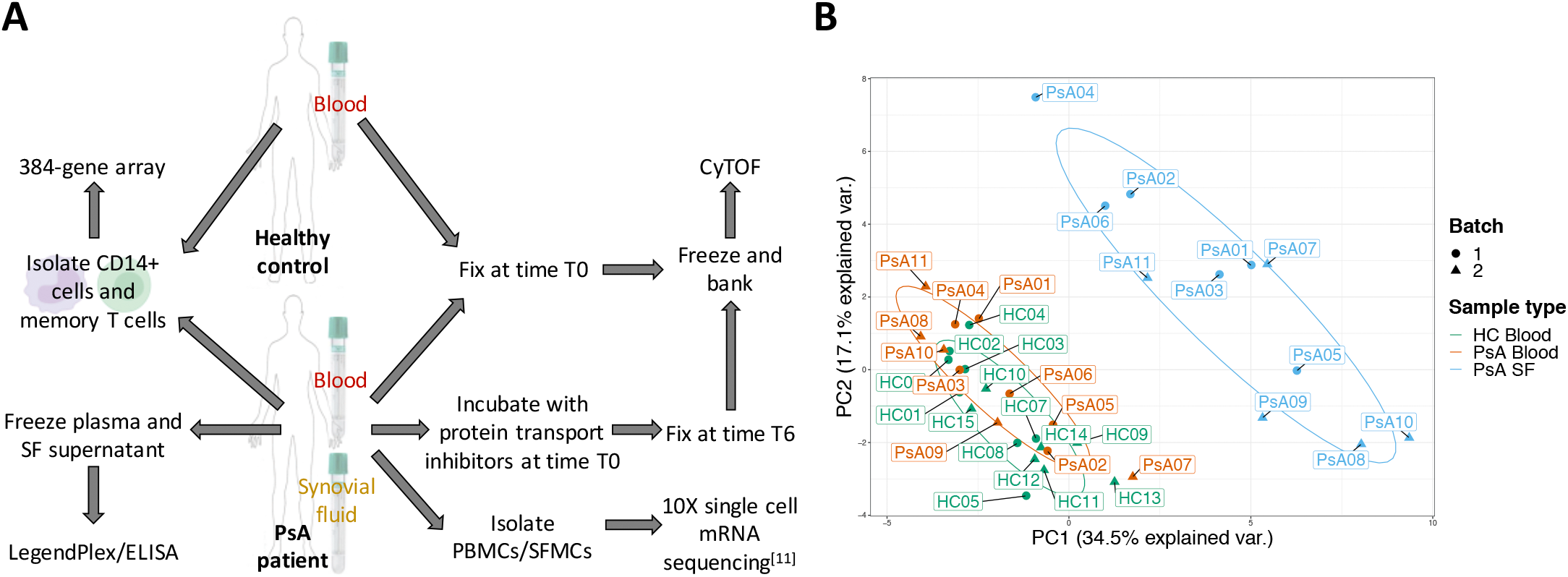
Overview of the experimental workflow and clear distinction of PsA synovial fluid (SF) and blood by principal component analysis. (A) Experimental workflow. Fresh peripheral blood and SF samples were split and either formaldehyde-fixed immediately following collection (time T0), or after incubation at 37°C with protein transport inhibitors for 6 h (time T6). These samples were used for phenotyping and intracellular CyTOF analysis. In addition, CD14+ cells, memory CD8+ T cells and memory CD4+ T cells were sorted and extracted RNA used in a 384-gene array; plasma and SF supernatant frozen; and PBMCs and SFMCs freshly isolated for 10X (previously described in [11]). (B) Unsupervised principal component analysis using the mean expression of lineage markers of CyTOF leucocyte samples at time T0 resolves PsA SF from matched and healthy control (HC) blood samples.

We next carried out an in-depth analysis using the unbiased clustering algorithm FlowSOM.^10^ Identified clusters were then manually annotated (supplementary table S2), merged and visualised using tSNE (figure 2A). Expression of key phenotypic markers of the different populations are represented as a heat map in figure 2B. The proportions of cell populations per patient are shown in figure 2C. We observed clear differences between the cell populations present in the SF compared to blood. Figure 2D shows that intermediate monocytes, macrophages, cDC1, cDC2, CD206+ cDC2, pDC, memory CD8 T cells, memory CD4 T cells and CD56bright NK cells were significantly increased in PsA SF (compared to PsA blood, all adjusted p-values < 0.01). Nonclassical monocytes, basophils, naïve CD8 T cells, naïve CD4 T cells, B cells, CD56dim CD16+ NK cells and NKT cells were significantly decreased in PsA SF (all adjusted p-values < 0.01).

Given that 4 of the 11 PsA patients were on methotrexate, we questioned whether this impacted cell population abundance. Methotrexate treatment did not affect monocyte or macrophage populations but increased cDC1, CD206+ cDC2 and CD56dim CD16+ NK cell populations, and reduced activated memory CD8 T cells (supplementary figure S1A). These methotrexate effects did not impact the overall increases/decreases seen in PsA SF compared to blood. We also compared healthy blood to PsA blood, finding a reduced frequency of pDC and MAIT cells in PsA blood (supplementary figure S1B).

**Figure 2.**
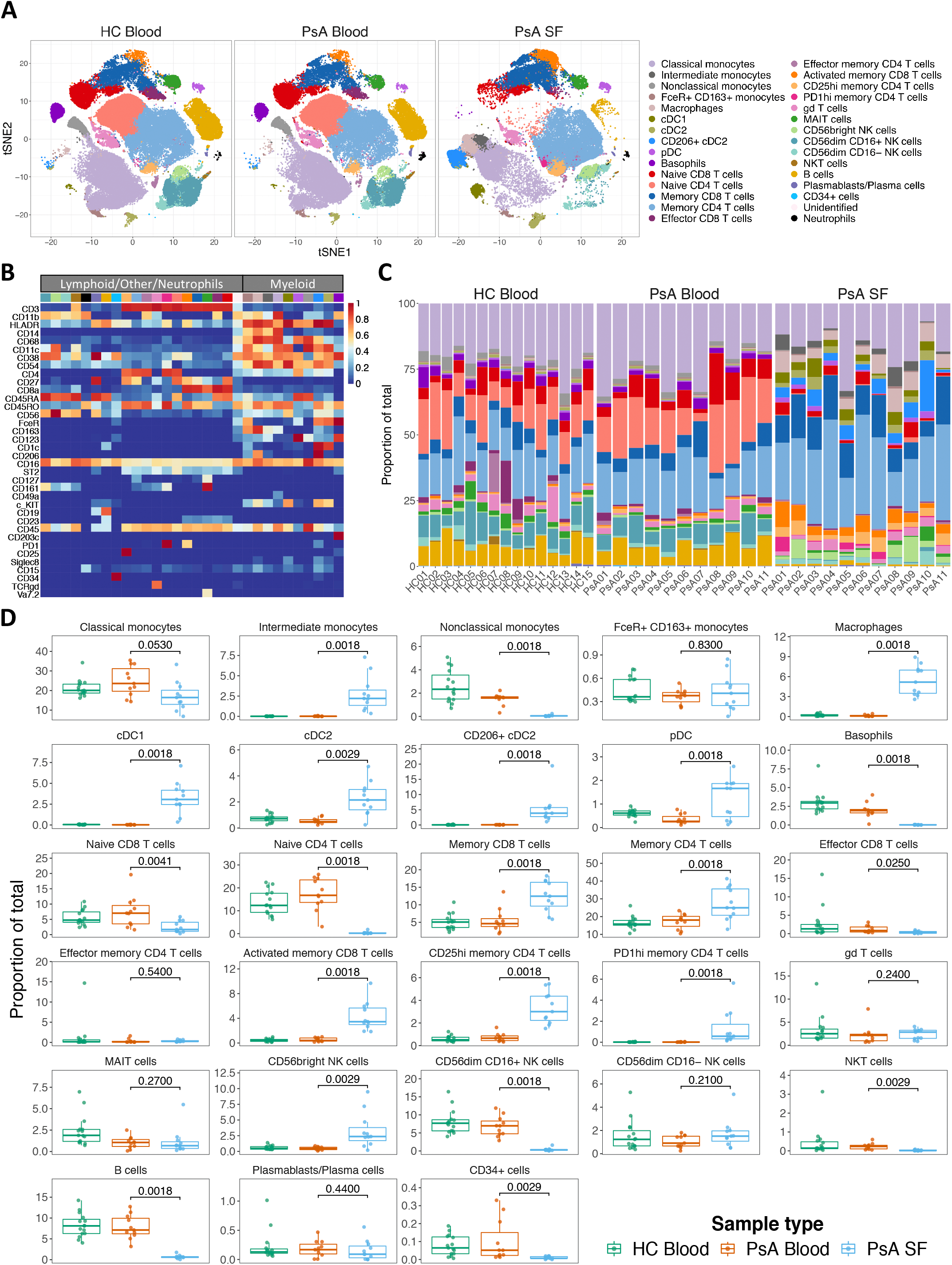
PsA SF CyTOF analysis shows expansion of multiple adaptive and innate cell populations compared to matched blood samples. (A) t-SNE plots showing leucocyte populations in 185,000 single cells (HC blood, n = 15; PsA blood, n = 11; PsA SF, n = 11, 5000 randomly selected cells from each sample). Cells are coloured according to the annotated and merged clusters, and stratified by sample type. (B) Heatmap of the median arcsinh-transformed marker intensity normalised to a 0 to 1 range of the 36 phenotyping panel markers across the 30 annotated clusters. (C) Cell composition for each individual studied stratified by sample type; neutrophils that were still present were omitted from this analysis. (D) Comparison of cluster frequencies in PsA blood and PsA SF. All p-values were calculated using paired Wilcox test and were corrected for multiple comparisons using the Benjamini-Hochberg adjustment at 5%. HC blood data is included for visualisation only. All samples were downsampled to an equal number of events (15,510 events) prior to clustering.

### Multiple memory T cell subsets are expanded in PsA SF

Given the expansion of memory T cells in PsA SF, we asked if any specific subpopulations were involved. Following data pre-processing and gating of CD3+ cells (supplementary figure S2A), FlowSOM was used to cluster the cell populations, which were then manually annotated and merged (supplementary figure S2B and C). PD1hi memory CD4 T cell, PD1mid memory CD4 T cell, CD25hi memory CD4 T cell and CD49a+ memory CD8 T cell populations were significantly increased in PsA SF compared to blood (supplementary figure S2D).

### PsA SF myeloid cells spontaneously release proinflammatory proteins on *ex vivo* incubation

To detect active protein production, we incubated matched SF and blood from 10 PsA patients for 6 h *ex vivo* (without any stimulation but in the presence of protein transport inhibitors to abrogate protein secretion) and compared these (T6) with matched T0 samples (figure 1A). In addition to 18 cell surface lineage markers, 18 intracellular markers were included to detect cytokines, chemokines and other secreted proteins, including IFNγ, IL-4, IL-10, IL-17 and IL-21 (predominantly secreted from T cells) and IL-8, CCL2, CXCL10, osteoactivin and osteopontin (predominantly myeloid). FlowSOM was again used to cluster the cell populations in an unsupervised manner followed by manual annotation of clusters (supplementary table S2), merging and visualisation using tSNE (figure 3A and supplementary figure S3A). In SF, osteopontin, CCL2 and IL-8 production were all significantly increased at T6 by at least 25% (with T6-T0 > 0.05%) in classical monocytes, intermediate monocytes and macrophages (mean expression osteopontin adjusted p-values = 1.37e-06, 1.99e-03 and 1.31e-04, respectively, CCL2 adjusted p-values = 3.31e-03, 1.34e-05 and 1.65e-04, respectively, and IL-8 adjusted p-values = 5.05e-04, 4.45e-08 and 2.83e-04, respectively) (figure 3B and supplementary table S3; other intracellular markers shown in supplementary figure S4). In addition, CXCL10 increased at T6 in intermediate monocytes when examining 95^th^ percentile expression (adjusted p-value = 6.32e-03) (supplementary figure S5).

In PsA blood, there were too few intermediate monocytes to analyse, however, in classical monocytes we observed at least a 25% increase in mean expression at T6 (with T6-T0 > 0.05%) in CCL2, IL8, IFNγ and IL-4 (adjusted p-value = 2.54e-09, 2.61e-05, 2.75e-07 and 5.33e-05) (supplementary figure S3B, S6 and supplementary table S3).

Following our unsupervised analysis, we reverted back to a manual biaxial analysis for visualisation and confirmation. Figure 3C shows production of osteopontin, CCL2 and IL-8 by PsA SF CD11c+ CD14+ CD123-cells over 6 h. Minimal osteopontin was detected in blood, while CCL2 was significantly higher in SF monocytes and IL-8 higher in blood monocytes (figure 3D). In addition, CXCL10 and osteoactivin production was higher in SF monocytes (supplementary figure S7).

**Figure 3.**
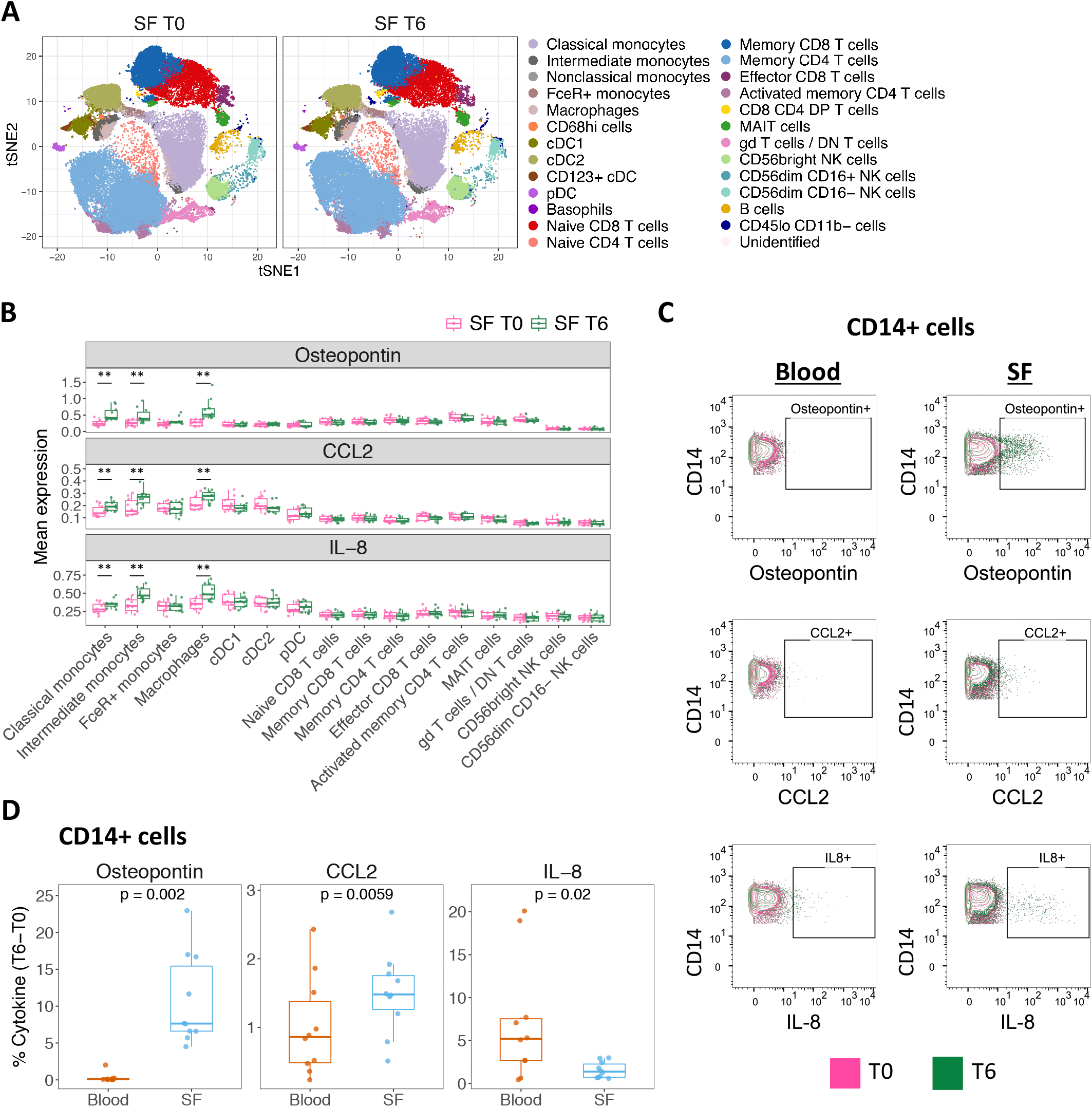
Osteopontin, CCL2 and IL-8 (CXCL8) are spontaneously produced by PsA SF monocytes/macrophages over 6 h *ex vivo*. (A) t-SNE plots based on the arcsinh-transformed expression of 18 markers in 5000 randomly selected cells from each sample (n = 10, only SF shown). Cells are coloured according to the annotated and merged clusters. Stratified by time. (B) Mean expression of osteopontin, CCL2 and IL-8 across the SF cell populations; any cell population containing <50 cells was omitted. ** indicates an overall increase in expression of at least 25% from T0 to T6 (with T6-T0 >0.05%), and a false discovery rate (FDR) <0.01. (C) Manual analysis of intracellular CyTOF data from a representative patient using FlowJo. Following data-preprocessing, .fcs files were gated on CD3-CD19-CD11c+ CD14+ CD123-cells. (D) Comparison of osteopontin, CCL2 and IL-8 production between SF and blood from manual analysis. The percentage of intracellular protein was calculated by subtracting the amount at time T0 from time T6 per patient sample for both blood and SF. All p-values were calculated using paired Wilcox test.

In terms of T cell cytokine release over the 6 h period, the only significant finding we identified was an increase in 95^th^ percentile expression of IL-10 in SF MAIT cells (adjusted p-value = 6.68e-03) (supplementary figure S5). A positive control using PMA/ionomycin stimulation was included in all three batches and demonstrated clearly the recall response of T cells and their ability to produce cytokines (supplementary figure S8).

### Gene expression analysis of SF T cells and monocytes shows upregulation of *SPP1* and *CCL2* compared to matched PBMCs

Next we sought to understand the *ex-vivo* transcriptomic signature of the key immune cell populations which differed in PsA SF compared to blood. Gene expression analysis of freshly cell sorted CD14+ cells, memory CD8 T cells and memory CD4 T cells from matched PBMC and SF samples (n = 3) was performed using a targeted array of 370 genes involved in inflammation or autoimmunity. As expected, the CD14+ cells clustered separately from the T cells across the PsA patients (figure 4A). For all three cell types, the majority of significantly dysregulated genes were upregulated in SF compared to blood (figure 4B). The gene for osteopontin, *SPP1*, was the highest upregulated gene (17.12 log2 fold change compared to blood) in SF CD14+ cells. *CCL2* and *CXCL10* were also upregulated in CD14+ cells, but not *CXCL8* (IL-8 gene). Despite significant upregulation of *OLR1* and *TNF*, their corresponding proteins were not significantly increased in the CyTOF dataset (figure 4C), although TNF protein was detected at time T6 in SF CD14+ cells in some patients (supplementary figure S7).

Gene expression in PsA PBMCs was also compared to three healthy controls (supplementary figure S9A). *SPP1* and *CCL2* are only upregulated in PsA SF vs PsA blood, with no significant difference when comparing PsA blood to healthy blood. *CXCL10* is upregulated in PsA SF but downregulated in PsA blood compared to healthy blood (supplementary figure S9B). To confirm the presence of secreted cytokines and chemokines in SF, PsA plasma and SF supernatant were analysed. Figure 4D shows that osteopontin, CCL2 and IL-8 were all enriched in SF compared to plasma. CXCL10 was previously shown to be increased in PsA SF.^11^

To confirm these findings we next interrogated single-cell RNA sequencing (scRNAseq) data from a recently described study of *ex vivo* PsA blood and SF.^11^ Unsupervised clustering of combined PBMC and SFMC data identified two clusters of monocytes/macrophages as defined by lineage markers *CD14, FCGR3A* (gene for CD16), *LYZ, MS4A7, CD163* and *MRC1* (gene for CD206); the smaller cluster was defined by increased expression of *APOE* (figure 5A). We observed strongest expression of *SPP1* and *CCL2* in the myeloid clusters (figure 5A). In this new analysis, we performed differential gene expression analysis between blood and SF in the monocyte/macrophage cluster and found *SPP1* to be the most selectively upregulated SF gene and *CCL2* the third most upregulated gene (adjusted p-values = 3.46e-239 and 8.08e-167, respectively, figure 5B). Both *SPP1* and *CCL2* were expressed by multiple cell populations in the SF, albeit at a lower level, including cDCs, NK cells and CD4 and CD8 T cells. Almost no expression of *SPP1* and *CCL2* was observed in the blood of any of the cell populations (figure 5C). For genes that had their corresponding proteins included in the CyTOF dataset, figure 5D shows that the monocyte/macrophage populations have the highest upregulation within the SF of a broad range of cytokine and chemokine genes.

**Figure 4.**
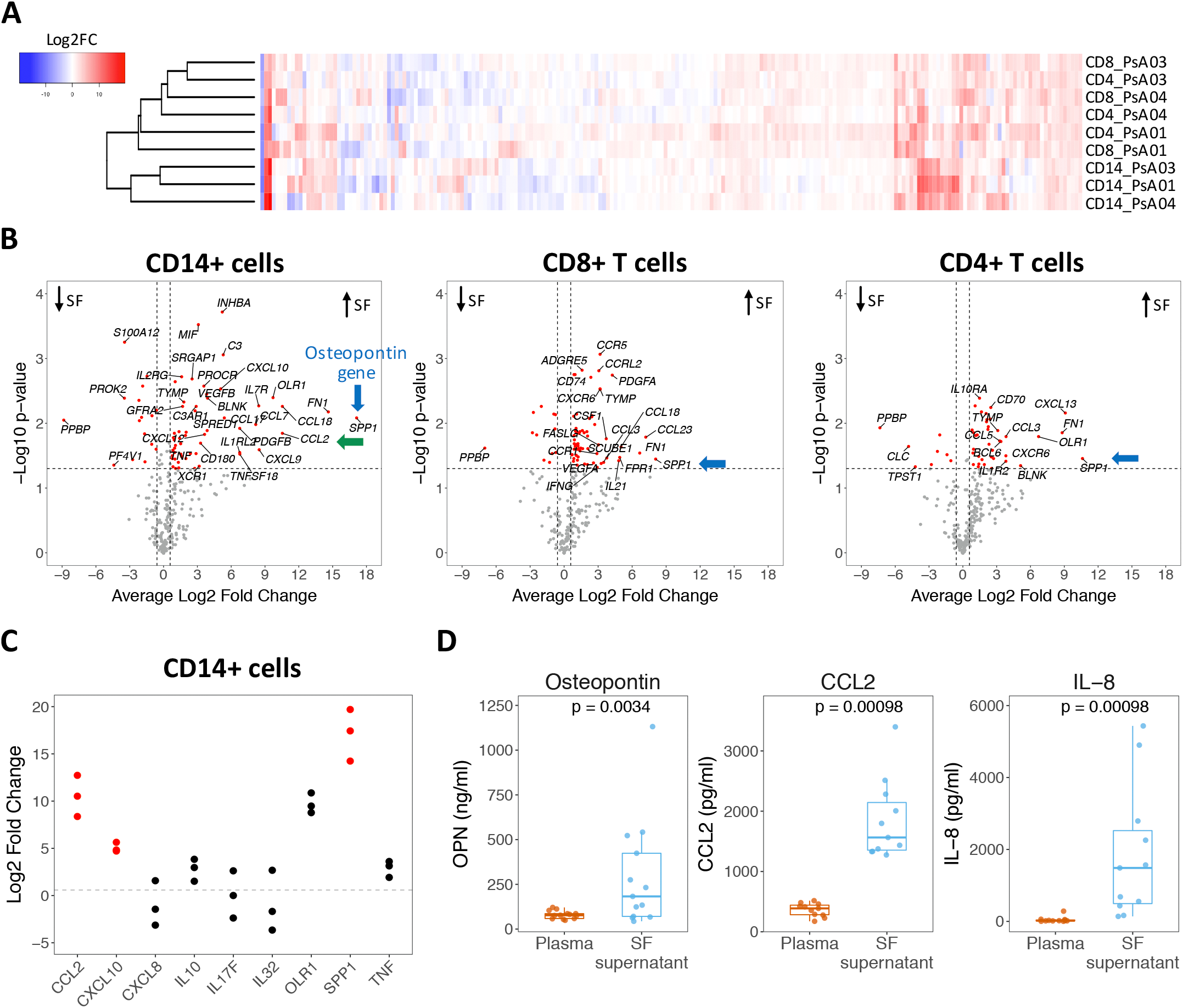
Gene expression analysis of isolated CD14+ cells, memory CD8+ T cells and memory CD4+ T cells from PsA SF compared to matched PBMCs (n = 3). (A) Heatmap of log2 gene expression fold change (FC) between SF and PBMCs for all genes that were detected. (B) Volcano plots showing differences in gene expression between SF and PBMC for CD14, CD8 and CD4 cells. The significance of the modulation in gene expression between the two tissues (y-axis) is plotted against the log2 of the mean FC (x-axis) across the three PsA patients. Genes showing p-value<0.05 (one sample t-test) and mean FC>1.5 are coloured in red. Black arrows indicate the direction of upregulation and downregulation of transcripts in SF; blue arrows point to *SPP1*, the gene for osteopontin, and green arrow points to *CCL2*. (C) Log2 FC in CD14+ cells for the genes that had their proteins included in the CyTOF intracellular panel. Genes that have corresponding proteins that were significantly increased in SF after 6 h as determined by CyTOF are coloured in red. (D) Osteopontin, CCL2 and IL-8 protein quantification in paired plasma and SF; CCL2 and IL-8 were measured by LegendPlex (n = 11); osteopontin was measured by ELISA (n = 13).

**Figure 5.**
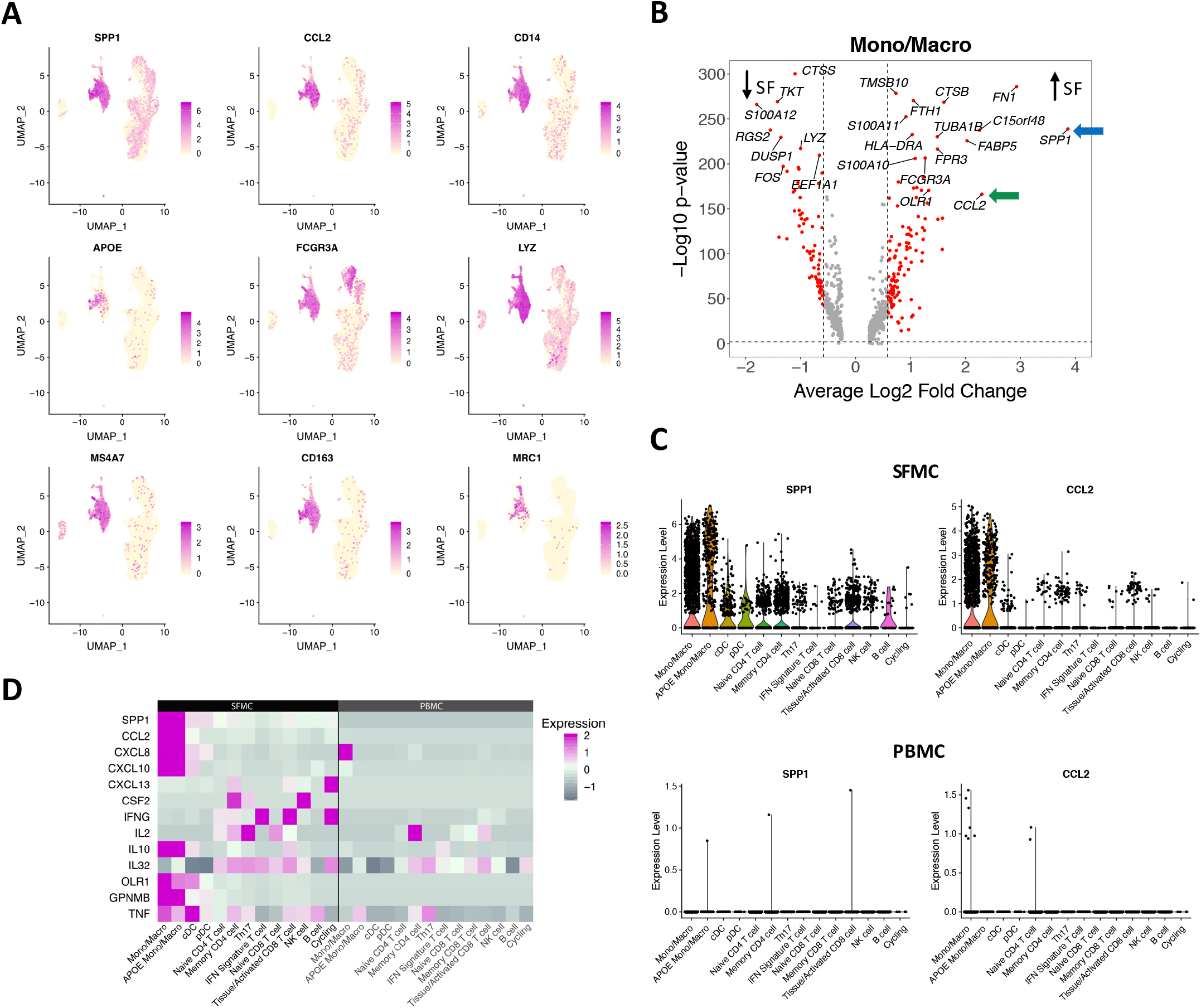
The genes for osteopontin and CCL2 are highly upregulated in monocytes/macrophages in a scRNA sequencing dataset of PsA SFMCs compared to matched PBMCs. (A) UMAPs of integrated PsA paired SFMC and PBMCs generated by 10× 3′ sequencing showing the relative expression of key annotated genes (n = 3). (B) Volcano plot showing differences in gene expression. The significance of the modulation in gene expression between the two tissues (y-axis) is plotted against the log2 of the mean FC (x-axis) across the three PsA patients. Genes showing corrected p-value<0.01 and mean FC>1.5 are coloured in red. Black arrows indicate the direction of upregulation and downregulation of transcripts in SF; blue arrow points to *SPP1*, the gene for osteopontin, and green arrow points to *CCL2*. (C) Violin plots of *SPP1* and *CCL2* across all clusters based on log normalised RNA. (D) Heatmap of gene expression across all clusters for genes that had their proteins included in the CyTOF intracellular panel; any gene that was not expressed in any cluster has been omitted.

## Discussion

In this study we use multiple complementary approaches to characterise the cellular and inflammatory landscape in PsA directly *ex vivo* and identify expansions of immunologically active myeloid populations within the joints. In order to minimise artefact and best capture the *in vivo* environment, we utilised fresh whole SF and blood^12^ for our CyTOF assays, with a 6 h incubation window to allow interrogation of the intrinsic cytokine/chemokine secretion profile.^13^ Here we show expansion of myeloid populations within the PsA joints which spontaneously produce proinflammatory cytokines and chemokines.

We observed a marked expansion of intermediate monocytes in PsA SF. These cells exhibit features in common with macrophages,^14^ are likely proinflammatory,^15^ and resemble those seen in rheumatoid arthritis (RA) and inflammatory osteoarthritis, where they correlate with disease activity.^16 17^ Unsupervised analysis identified spontaneous high levels of production of osteopontin, CCL2 and IL-8 by SF intermediate monocytes, classical monocytes and macrophages. CXCL10 production by SF intermediate monocytes was also observed. CD14+ cells in SF produced significantly more osteopontin, CCL2 and CXCL10 than blood, and our findings indicate that they are almost certainly the main cellular source of these factors which are enriched in the SF.

We were able to confirm our protein findings at the gene level, with *SPP1* (the gene for osteopontin) being the highest upregulated gene both in freshly sorted PsA SF CD14 monocytes/macrophages (compared to PsA blood CD14+ cells) and in synovial monocytes/macrophages in a large scRNAseq dataset. *CCL2* and *CXCL10* upregulation also matched protein expression in myeloid populations. Interestingly osteopontin was not detected by CyTOF in SF T cells, nor CXCL13 in SF CD4 T cells, despite their genes being significantly upregulated, emphasising the importance of quantifying protein expression.^18^ When comparing PsA SF and blood to healthy blood, *CXCL10, TNF* and *CCL5* were upregulated in PsA SF CD14+ cells but downregulated in PsA blood compared to healthy blood, indicating potential trafficking of these cells into the SF.

Our study suggests a potential important role for myeloid cell production of osteopontin in PsA pathogenesis. Osteopontin showed the greatest increase in PsA SF CD14 cells in independent bulk and scRNAseq datasets, was spontaneously produced by PsA SF (not blood) myeloid populations and was present at increased levels in PsA SF. Osteopontin induces chemotactic migration of both macrophages and T cells,^19 20^ stimulates Th1 and Th17 cytokine release and downmodulates IL-10.^21-24^ Previous work has shown that osteopontin serum and SF levels correlate with CRP in RA patients.^25 26^ Although we did not observe this correlation in our PsA patients (data not shown), a larger patient cohort would be required to form a conclusion. In addition, osteopontin has been shown to induce the expression of CCL2 and CCL4 in monocytes.^25^ In both our CyTOF and gene expression assays, osteopontin and CCL2 were increased in the monocyte/macrophage population, demonstrating they may be intrinsically linked. Osteopontin has been reported as upregulated in PsA synovial biopsies,^27^ and here we show that it is highly expressed in PsA SF, with by far the greatest production from the myeloid compartment. In patients with RA, serum levels of osteopontin predict effectiveness of tocilizumab,^28^ although osteopontin neutralization failed to induce clinical improvement,^29^ perhaps due to rapid turnover.^30^

Both CXCL10 and CCL2 have been reported as upregulated in PsA serum^31^ with CXCL10 increased in PsA SF^32^ and CCL2 upregulated in RA SF.^33^ We have previously suggested a potential role for the CXCL10 receptor CXCR3 on CD8 T cells in PsA pathogenesis,^11^ and here we show that PsA SF monocytes are a key cellular source of CXCL10, likely contributing to the recruitment of these cells. A clinical trial of RA patients that combined a monoclonal antibody targeting CXCL10 with methotrexate had a modest clinical effect.^34^ The best described function of CCL2 is monocyte recruitment,^35^ however, it has pleiotropic effects on myeloid cells^36^ and is capable of recruiting other cell types including T cells.^37^ Therefore it may play a central role in both myeloid and T cell recruitment in PsA and may represent an opportunity to intervene therapeutically. CCL2 inhibition has shown efficacy in rat adjuvant arthritis,^38^ but not RA.^39^ CCL2 inhibition has not been tested in PsA, and our data would support such study.

Although we detected IL-8 (CXCL8) production by PsA monocytes/macrophages this was greater in blood than SF and it is likely that synovial tissue cells or neutrophils are the dominant source for this cytokine in PsA joints. Our study did not look at either of these cell types and this will be important to examine in future.

Our CyTOF approach allowed detailed study of both myeloid and lymphoid populations present in PsA SF and blood. We here confirm previous findings where technical considerations frequently only allowed focus on a particular cell type.^8 11 40-42^ We observed SF expansion of PD1hi memory CD4 T cells, representing T follicular helper cells/peripheral T helper cells, similar to that seen in rheumatoid arthritis (RA),^43 44^ and of tissue-resident CD49a+ memory CD8 T cells^6^ that we previously showed to be clonally expanded in PsA.^11^ Interestingly these cells are phenotypically similar to the integrin-expressing (InEx) cells recently described in the joints in related SpA ankylosing spondylitis.^45 46^ Although PsA SF T cells have the capability to produce cytokines such as IFNγ, IL-17 and GM-CSF upon *in vitro* stimulation,^47^ we did not see any significant T cell cytokine production in our *ex vivo* unstimulated CyTOF assay. The ability to capture the exact moment when these cells are stimulated *in vivo* and exit dormancy may require a longer period of incubation or may be beyond current detection capabilities.

In summary, we have used direct *ex vivo* CyTOF analysis, validated by gene expression analysis to identify expanded SF monocytes/macrophage populations that are actively and spontaneously producing cytokines. These may be of diagnostic, prognostic and/or therapeutic importance.

## Methods

### Study subjects and patient involvement

The study was conducted in accordance with protocols approved by the Oxford Research Ethics committee (Ethics reference number 06/Q1606/139). Patients with PsA were involved throughout in discussions about areas of research but not in detailed study design. Blood and SF samples were collected from 11 consecutive patients not receiving biologic DMARDS or steroids (6 males, 5 females, mean age 43.8 ± 13.5 years; 4 patients on methotrexate) with large-joint oligo PsA undergoing intra-articular knee aspiration at Oxford University Hospitals; 3 of these were assessed by PCR array and 3 for scRNA sequencing. An additional 3 patients were included for targeted SF and plasma protein analysis. Blood from 15 anonymous healthy donors (10 males, 5 females, mean age 45.1 ± 10.4 years) was collected under UCB Celltech UK HTA license number 12504. Full informed consent was obtained from all subjects. Demographics for all study subjects are listed in supplementary table S1.

### CyTOF staining and analysis

Fresh SF and blood were split and either immediately (within 30 minutes of sampling) fixed with high-purity paraformaldehyde (time T0) or incubated at 37°C, 5% CO_2_ at a 1:1 ratio with RPMI 1640 containing brefeldin A and monensin for 6 h (time T6) with Intercalator-103Rh added 15 min prior to fixation. All samples were then stored at -80°C. Samples were lysed with Permeabilization Buffer (eBioscience), barcoded and cells stained in Maxpar staining buffer (Fluidigm) with antibodies listed in supplementary table S4. Samples were run on a Helios instrument alongside normalization beads (Fluidigm). Raw CyTOF files were debarcoded using Fluidigm software, and the .fcs files were pre-processed in FlowJo using biaxial manual gating as shown in supplementary figure S10. These pre-processed files were analysed using a modified custom R workflow^48^ with cell populations clustered using ConsensusClusterPlus^49^ and FlowSOM algorithms^10^ – a highly recommended clustering method.^50^ Clusters were merged and annotated as defined in supplementary table S2.

### Quality control of CyTOF assays and analysis

Samples were processed in two batches for the phenotyping panel and three batches for the intracellular panel (supplementary table S1). The following strategies were undertaken to minimise batch effects and bias: barcoding of samples; thawing and staining of samples on the same day with the same antibody mix; inclusion of a technical control (supplementary figures S11 and S12); and unsupervised clustering of data using multiple seeds (supplementary figure S13). Paired samples were always kept in the same batch. Whilst we detected a batch effect for macrophages (and CD34+ cells), in both batches macrophages were increased in PsA SF compared to blood (supplementary figure S1C). For intracellular panel analyses, we set a minimum increase in protein production of 25% and a difference in T0 from T6 of at least 0.05%. Differential analysis of marker expression was conducted using a linear mixed model with time-point and batch as fixed effects, and the paired patient sample as a random effect. Multiple comparisons were corrected using the Benjamini-Hochberg adjustment.

### Cell isolation and sorting

PBMC and SFMCs were isolated from venous blood by density centrifugation (Histopaque; Sigma). PBMCs and SFMCs were stained with the following antibodies: live/dead eFluor780; CD14 PE-Cy7 (M5E2); CD3 FITC (SK7); CD4 APC (RPA-T4); CD8a PE (RPA-T8); CD45RA BV421 (HI100) (all BioLegend). Live CD3-CD14+, CD3+CD8+CD45RA- and CD3+CD4+CD45RA-cell populations were bulk sorted then frozen into RLT buffer (Qiagen).

### PCR array

RNA was isolated using the RNeasy Micro kit (Qiagen), and a custom 384-gene array (RT^2^ Profiler PCR array; Qiagen) focused on the inflammatory response/autoimmunity was performed following the manufacturer’s instructions. Relative fold expression levels were calculated after normalisation against the housekeeping genes, then using the ΔΔCt method to calculate normalised gene expression in the SF compared to matched blood sample. In addition, PsA blood samples were compared to the mean expression of three healthy donors. Genes that were undetected or with a threshold cycle >35 were omitted from further analyses. Genes were considered significant using a p-value <0.05 (one sample t-test) and a mean fold change >1.5.

### Protein quantitation by multiplex assay and Enzyme linked immunosorbent assay (ELISA)

Matched plasma and SF supernatants were collected and stored at -80°C. Samples were thawed and proteins quantified using a LEGENDplex Human Proinflammatory Chemokine Panel immunoassay (BioLegend) according to the manufacturer’s instructions. Data were acquired on a Novocyte instrument and analysed using software provided by BioLegend. To quantify osteopontin, the Human Osteopontin ELISA kit (Thermo Fisher) was used following the manufacturer’s instructions. All p-values were calculated using paired Wilcox test.

### scRNA sequencing analysis

We performed further analysis of the single cell data from a previously published dataset.^11^ All pre-processing, quality control, alignment and basic clustering of combined PBMC and SFMC samples was carried out as previously published. In this analysis, we used Seurat (Satija lab, version 3.0) to split the clusters according to their tissue of origin (PBMC vs SFMC) based on the metadata and performed differential gene expression analysis for the two main myeloid clusters. We also plotted the expression of *SPP1* and *CCL2* across all clusters in the split PBMC/SFMC analysis and used the average expression analysis function to generate the heatmap of chemokine/cytokine expression split across PBMC and SFMC.

### Statistical analysis

All statistical analyses were performed using R. Statistical tests were used as stated in the figure legends.

## Acknowledgements and funding

At UCB, we thank Catherine Simpson for assistance with the cell sorting and CyTOF sample acquisition, and at Idorsia Pharmaceuticals, we thank Andrew Croxford for assistance with the LEGENDplex assay. NY and HAM received funding from UCB and HAM from National Institute for Health Research (NIHR). SC and AM are employees of UCB. PB has received research support from Regeneron, Benevolent AI and GSK. The study received support from the NIHR Oxford Biomedical Research Centre (BRC) (PB). The views expressed are those of the author(s) and not necessarily those of the NHS, the NIHR or the Department of Health.

