## Supplementary Material for "*Ex vivo* mass cytometry analysis reveals a profound myeloid proinflammatory signature in psoriatic arthritis synovial fluid"

### Supplementary Table S1

#### Samples for CyTOF

| Donor ID | Gender | Age | Treatment | CRP (mg/L) | Phenotyping Batch | Intracellular Batch |
|---|---|---|---|---|---|---|
| HC01 | M | 57 | | | 1 | nd |
| HC02 | F | 61 | | | 1 | nd |
| HC03 | M | 45 | | | 1 | nd |
| HC04 | M | 47 | | | 1 | nd |
| HC05 | F | 47 | | | 1 | nd |
| HC06* | F | 43 | | | 1 and 2 | 1, 2 and 3 |
| HC07 | M | 53 | | | 1 | nd |
| HC08 | M | 52 | | | 1 | nd |
| HC09 | F | 46 | | | 2 | nd |
| HC10 | F | 32 | | | 2 | nd |
| HC11 | M | 38 | | | 2 | nd |
| HC12 | M | 31 | | | 2 | nd |
| HC13 | M | 60 | | | 2 | nd |
| HC14 | M | 33 | | | 2 | nd |
| HC15 | M | 31 | | | 2 | nd |
| PsA01 | F | 72 | None | 98.6 | 1 | 1 |
| PsA02 | M | 61 | None | 33.8 | 1 | 1 |
| PsA03 | M | 34 | MTX | 15.8 | 1 | 1 |
| PsA04 | M | 42 | None | 8.0 | 1 | 1 |
| PsA05 | M | 49 | MTX | 58.7 | 1 | 2 |
| PsA06 | F | 31 | None | 12.5 | 1 | 2 |
| PsA07 | F | 53 | None | 9.9 | 2 | 2 |
| PsA08 | M | 35 | None | 1.0 | 2 | 2 |
| PsA09 | F | 33 | MTX | 16.2 | 2 | 3 |
| PsA10 | M | 41 | MTX | 9.0 | 2 | 3 |
| PsA11 | F | 31 | None | nd | 2 | nd |

#### Samples for PCR array

| Donor ID | Gender | Age | Treatment | CRP (mg/L) |
|---|---|---|---|---|
| PsA01 | F | 72 | None | 43.2 |
| PsA03 | M | 33 | None | 36.6 |
| PsA04 | M | 42 | None | 8.0 |

#### Samples for 10X

| Donor ID | Gender | Age | Treatment | CRP (mg/L) |
|---|---|---|---|---|
| PsA04 | M | 42 | None | 8.0 |
| PsA07 | F | 53 | None | 9.9 |
| PsA08 | M | 35 | None | 1.0 |

#### Samples for LEGENDplex/ELISA

| Donor ID | Gender | Age | Treatment | CRP (mg/L) | LEGENDplex/ELISA |
|---|---|---|---|---|---|
| PsA01 | F | 72 | None | 43.2 | Both |
| PsA02 | M | 61 | None | 33.8 | Both |
| PsA03 | M | 33 | None | 36.6 | LEGENDplex |
| PsA04 | M | 42 | None | 8 | Both |
| PsA05 | M | 49 | MTX | 58.7 | Both |
| PsA06 | F | 31 | None | 25.4 | Both |
| PsA07 | F | 53 | None | 9.9 | Both |
| PsA08 | M | 35 | None | 1 | Both |
| PsA09 | F | 33 | MTX | 16.2 | Both |
| PsA10 | M | 41 | MTX | 9.0 | ELISA |
| PsA11 | F | 31 | None | nd | ELISA |
| PsA12 | F | 26 | None | 1.8 | Both |
| PsA13 | M | 65 | Leflunomide, Etanercept | 7.2 | Both |
| PsA14 | F | 28 | None | 0.04 | ELISA |

**Supplementary Table S1. Demographics of PsA patients and controls studied, including staining batch for CyTOF.**
Matched SF and blood were included in the same batch for CyTOF for all PsA patients.
*One healthy control (HC06) was included in all CyTOF batches as a technical control.

#### Supplementary Table S2

| Cluster name | Phenotyping panel | Intracellular panel |
|---|---|---|
| Naive CD8 T cells | CD45+ CD3+ CD8a+ CD45RA+ CD27+ CD4- CD56- | CD45+ CD3+ CD8a+ CD45RO- CD4- CD56- |
| Naive CD4 T cells | CD45+ CD3+ CD4+ CD45RA+ CD27+ CD8a- CD56- | CD45+ CD3+ CD4+ CD45RO- CD8a- CD56- |
| Memory CD8 T cells | CD45+ CD3+ CD8a+ CD4- CD45RO+ CD56- | CD45+ CD3+ CD8a+ CD4- CD45RO+ CD56- |
| Memory CD4 T cells | CD45+ CD3+ CD4+ CD8a- CD45RO+ CD27+ CD56- | CD45+ CD3+ CD4+ CD8a- CD45RO+ CD56- |
| Activated memory CD8 T cells | CD45+ CD3+ CD8a+ CD4- CD45RO+ HLA-DR+ CD38+ PD1+ CD56- | n/a |
| Effector CD8 T cells | CD45+ CD3+ CD8a+ CD45RA+ CD11b_mid CD27- CD4- CD56- | CD45+ CD3+ CD8a+ CD45RO- CD11b_mid CD4- CD56- |
| Effector memory CD4 T cells | CD45+ CD3+ CD4+ CD45RO+ CD11b_mid CD27- CD8a- CD56- | n/a |
| Activated memory CD4 T cells | n/a | CD45+ CD3+ CD4+ CD45RO+ HLA-DR+ CD8a- CD56- |
| CD8 CD4 DP T cells | n/a | CD45+ CD3+ CD8a+ CD4+ CD56- |
| PD1hi memory CD4 T cells | CD45+ CD3+ CD4+ CD8a- CD45RO+ HLA-DR+ PD1hi CD56- | n/a |
| CD25hi memory CD4 T cells | CD45+ CD3+ CD4+ CD8a- CD45RO+ CD27+ HLA-DR+ CD25hi CD56- | n/a |
| Classical monocytes | CD45+ CD11b+ CD14+ CD11c+ CD68+ HLA-DR+ CD163+ CD4lo | CD45+ CD11b+ CD14+ CD11c+ CD68+ HLA-DR+ CD4lo |
| Intermediate monocytes | CD45+ CD14lo CD16mid CD11c+ HLA-DR+ CD11b_mid CD68+ CD163+ CD123lo | CD45+ CD14lo CD16mid CD11c+ HLA-DR+ CD11b_mid CD68+ CD123lo |
| Nonclassical monocytes | CD45+ CD16hi CD11c+ CD68+ CD45RA+ HLA-DR+ CD11b_lo CD38lo CD123lo CD14- CD163- | CD45hi CD16+ CD11c+ CD68+ CD45RO- HLA-DR+ CD11b_lo CD123lo CD14- |
| FceR+ CD163+ monocytes | CD45+ CD11b+ CD14+ CD11c+ CD68+ HLA-DR+ FceR+ CD163+ CD4lo | n/a |
| FceR+ monocytes | n/a | CD45+ CD11b+ CD14+ CD11c+ CD68+ HLA-DR+ FceR+ CD4lo |
| Macrophages | CD45+ CD11b+ HLA-DR+ CD14mid CD68+ CD11c+ CD4lo CD56lo CD163hi CD206+ | CD45+ CD11b+ HLA-DR+ CD14mid CD68+ CD11c+ CD4lo CD56lo |
| B cells | CD45+ CD19+ HLA-DR+ CD45RA+ | CD45+ CD19+ HLA-DR+ CD45RO- |
| Plasmablasts/Plasma cells | CD45+ CD19+ HLA-DR+ CD45RA+ CD38hi CDD27+ | n/a |
| Basophils | CD45lo CD11b+ CD203c+ CD123+ FceR+ HLA-DR- | CD45lo CD11b+ CD123+ FceR+ HLA-DR- |
| gd T cells | CD45+ CD3+ TCRgd+ CD8a- CD4- CD56- | n/a |
| gd T cells / DN T cells | n/a | CD45+ CD3+ CD8a- CD4- CD56- |
| MAIT cells | CD45+ CD3+ CD8a+ CD161hi Va7.2+ CD56- | CD45+ CD3+ CD8a+ CD161hi CD56- |
| CD56bright NK cells | CD45+ CD56hi CD11b+ CD11c+ CD38+ CD45RA+ CD161+ CD3- | CD45+ CD56hi CD11b+ CD11c+ CD45RO- CD161+ CD3- |
| CD56dim CD16+ NK cells | CD45+ CD56+ CD16+ CD11b+ CD11c+ CD38+ CD45RA+ CD161+ CD3- | CD45+ CD56+ CD16+ CD11b+ CD11c+ CD45RO- CD161+ CD3- |
| CD56dim CD16- NK cells | CD45+ CD56+ CD11b+ CD11c- CD45RA+ CD161+ CD3- | CD45+ CD56+ CD11b+ CD11c- CD45RO- CD161+ CD3- |
| NKT cells | CD45+ CD3+ CD56+ CD11b+ | n/a |
| pDC | CD45lo CD123hi CD68+ HLA-DR+ CD45RA+ CD4+ CD14- CD11b- CD11c- CD56lo | CD45lo CD123hi CD68+ HLA-DR+ CD45RO- CD4+ CD14- CD11b- CD11c- CD56lo |
| cDC1 | CD45+ CD11b- HLA-DR+ CD14- CD68lo CD11c+ CD4lo FceR- CD123- CD1c- CD206- | CD45+ CD11b- HLA-DR+ CD14- CD68lo CD11c+ CD4lo FceR- CD123- |
| cDC2 | CD45+ CD11c+ CD1c+ FceR+ HLA-DR+ CD68lo CD56lo CD4lo CD11b_lo CD14- CD123- CD206- | CD45+ CD11c+ FceR+ HLA-DR+ CD68lo CD56lo CD4lo CD11b_lo CD14- CD123- |
| CD206+ cDC2 | CD45+ CD11c+ CD1c+ FceR+ HLA-DR+ CD68lo CD56lo CD4lo CD11b_lo CD14- CD123- CD206+ | n/a |
| CD123+ cDC | n/a | CD45+ CD11b- HLA-DR+ CD14- CD68lo CD11c+ CD4lo FceR- CD123+ |
| CD34+ cells | CD45lo CD34+ CD38+ HLA-DR+ | n/a |
| CD68hi cells | n/a | CD45+ CD68hi HLA-DR+ CD14- CD11b_lo CD11c_lo CD4lo CD56lo |
| CD45lo CD11b- cells | n/a | CD45lo CD11b- CD11c- CD45RO- |
| Neutrophils | CD45lo CD11b+ CD15+ CD16+ | CD45lo CD11b+ CD15+ CD16+ |

**Supplementary Table S2. Defining CyTOF markers used for cell population clusters.** FlowSOM and ConsensusClusterPlus algorthims were used to generate 40 clusters for the phenotyping panel and 35 clusters for the intracellular panel. A heatmap of these clusters led to final cluster annotation and merging.

### Supplementary Table S3

## SF

| Cell population | Protein | Median SF T0 | Median SF T6 | Median SF T6-T0 | Median SF T6/T0 | Adjusted p-value T0vsT6 |
|---|---|---|---|---|---|---|
| Macrophages | Osteopontin | 0.277827984 | 0.514038861 | 0.236210878 | 1.850205493 | 0.00013116 |
| Intermediate monocytes | CCL2 | 0.154485682 | 0.27235715 | 0.117871468 | 1.762992828 | 1.33733E-05 |
| Classical monocytes | Osteopontin | 0.23731464 | 0.413602648 | 0.176288008 | 1.742845058 | 1.37057E-06 |
| Intermediate monocytes | Osteopontin | 0.258648552 | 0.38388891 | 0.125240358 | 1.484210551 | 0.001985722 |
| Intermediate monocytes | IL-8 | 0.322100649 | 0.466827381 | 0.144726732 | 1.449321455 | 4.45362E-08 |
| Classical monocytes | CCL2 | 0.136908377 | 0.191444846 | 0.054536469 | 1.398342819 | 0.003305701 |
| Macrophages | CCL2 | 0.202538038 | 0.279908058 | 0.07737002 | 1.382002419 | 0.000164875 |
| Macrophages | IL-8 | 0.351329244 | 0.482511672 | 0.131182428 | 1.373388865 | 0.000283155 |
| Activated memory CD4 T cells | IL-10 | 0.060697847 | 0.080484555 | 0.019786708 | 1.325986988 | 0.003567156 |
| Classical monocytes | IL-8 | 0.274873037 | 0.352231238 | 0.077358201 | 1.28143248 | 0.000505323 |

#### Blood

| Cell population | Protein | Median SF T0 | Median SF T6 | Median SF T6-T0 | Median SF T6/T0 | Adjusted p-value T0vsT6 |
|---|---|---|---|---|---|---|
| Classical monocytes | IL-8 | 0.188306715 | 0.558024088 | 0.369717373 | 2.963378592 | 2.60837E-05 |
| Classical monocytes | CCL2 | 0.091330313 | 0.149204418 | 0.057874105 | 1.633679039 | 2.54237E-09 |
| Classical monocytes | IFNγ | 0.106513338 | 0.158724792 | 0.052211454 | 1.490187004 | 2.75422E-07 |
| Classical monocytes | IL-4 | 0.154039676 | 0.226332765 | 0.072293089 | 1.46931473 | 5.32833E-05 |
| Classical monocytes | Osteopontin | 0.115721607 | 0.156416819 | 0.040695212 | 1.351664767 | 0.000540984 |
| Classical monocytes | CXCL10 | 0.161304402 | 0.207236918 | 0.045932516 | 1.284756742 | 5.57158E-05 |
| Classical monocytes | CXCL13 | 0.112883992 | 0.144937006 | 0.032053014 | 1.283946496 | 2.60837E-05 |

**Supplementary Table S3. Adjusted p-values for intracellular proteins within cell populations shown in Fig 3 and Supp Fig S3.** Only significant protein production in cell populations that have a T6/T0 value > 1.25 are shown. Differential analysis of marker expression stratified by cell population was conducted using a linear mixed model whereby time-point and batch were fixed effects, and the paired patient sample was a random effect. Multiple comparisons were corrected using the Benjamini-Hochberg adjustment.

#### Supplementary Table S4

| | Phenotyping panel | | | | Intracellular panel | | | |
|---|---|---|---|---|---|---|---|---|
| Metal | Target | Clone | Supplier | Used for cell clustering | Target | Clone | Supplier | Used for cell clustering |
| 89Y | CD45 | HI30 | Fluidigm | Yes | CD45 | HI30 | Fluidigm | Yes |
| 103Rh | | | | | Live/dead | | Fluidigm | No |
| 141Pr | Siglec-8 | 7C9 | BioLegend | Yes | OLR1 | 331212 | R&D Systems | No |
| 142Nd | CD19 | HIB19 | Thermo Fisher (eBioscience) | Yes | CD19 | HIB19 | Thermo Fisher (eBioscience) | Yes |
| 143Nd | Va7.2 | 3C10 | BioLegend | Yes | CCL2 | 5D3-F7 | BioLegend | No |
| 144Nd | CD15 | W6D3 | BioLegend | Yes | CXCL13 | 53610 | R&D Systems | No |
| 145Nd | CD4 | RPA-T4 | Thermo Fisher (eBioscience) | Yes | CD4 | RPA-T4 | Thermo Fisher (eBioscience) | Yes |
| 146Nd | CD8a | RP4-T8 | Thermo Fisher (eBioscience) | Yes | CD8a | RP4-T8 | Thermo Fisher (eBioscience) | Yes |
| 147Sm | CD34 | 4H11 | Thermo Fisher (eBioscience) | Yes | IL8 | G265-8 | BD Biosciences | No |
| 148Nd | CD16 | 3G8 | Thermo Fisher (eBioscience) | Yes | CD16 | 3G8 | Thermo Fisher (eBioscience) | Yes |
| 149Sm | CD25 | BC96 | Thermo Fisher (eBioscience) | Yes | | | | |
| 150Nd | CD163 | GHI/61 | BioLegend | Yes | IL4 | MP4-25D2 | BioLegend | No |
| 151Eu | CD123 | 6H6 | Thermo Fisher (eBioscience) | Yes | CD123 | 6H6 | Thermo Fisher (eBioscience) | Yes |
| 152Sm | PD1 | EH12.2H7 | BioLegend | Yes | IL21 | 3A3-N2 | BioLegend | No |
| 153Eu | CD1c | L161 | BioLegend | Yes | IL17F | SHLR17 | BioLegend | No |
| 154Sm | CD206 | 15-2 | BioLegend | Yes | IL2 | MQ1-17H12 | Thermo Fisher (eBioscience) | No |
| 155Gd | | | | | TNFa | MAb1 | Thermo Fisher (eBioscience) | No |
| 156Gd | CD203c | NP4D6 | Thermo Fisher (eBioscience) | Yes | IL17A | eBio64CAP17 | Thermo Fisher (eBioscience) | No |
| 158Gd | CD49a | TS2/7 | BioLegend | Yes | IL10 | JES3-9D7 | Thermo Fisher (eBioscience) | No |
| 159Tb | CD68 | FA-11 | BioLegend | Yes | CD11c | 3.9 | BioLegend | Yes |
| 160Gd | CD14 | 61D3 | Thermo Fisher (eBioscience) | Yes | CD14 | 61D3 | Thermo Fisher (eBioscience) | Yes |
| 161Dy | CD161 | DX12 | BD Biosciences | Yes | CD161 | DX12 | BD Biosciences | Yes |
| 162Dy | CD23 | M-L233 | BD Biosciences | Yes | IL6 | MQ2-13A5 | Thermo Fisher (eBioscience) | No |
| 163Dy | CD127 | A019D5 | BioLegend | Yes | IFNg | 45-15 | Miltenyi | No |
| 164Dy | TCRgd | B1/35 | BioLegend | Yes | GMCSF | BVD2-21C11 | BioLegend | No |
| 165Ho | FceR | AER-37 (CRA-1) | BioLegend | Yes | FceR | AER-37 (CRA-1) | BioLegend | Yes |
| 166Er | CD11c | 3.9 | BioLegend | Yes | CD15 | W6D3 | BioLegend | Yes |
| 167Er | CD27 | O323 | Thermo Fisher (eBioscience) | Yes | Osteopontin | 2F10 | Thermo Fisher (eBioscience) | No |
| 168Er | ST2 | polyclonal | R&D Systems | Yes | CXCL10 | 33036 | R&D Systems | No |
| 169Tm | CD45RA | HI100 | Thermo Fisher (eBioscience) | Yes | Osteoactivin | 303822 | R&D Systems | No |
| 170Er | CD3 | UCHT1 | Thermo Fisher (eBioscience) | Yes | CD3 | UCHT1 | Thermo Fisher (eBioscience) | Yes |
| 171Yb | CD45RO | UCHL1 | BioLegend | Yes | CD45RO | UCHL1 | BioLegend | Yes |
| 172Yb | CD38 | HIT2 | Thermo Fisher (eBioscience) | Yes | CD68 | FA-11 | BioLegend | Yes |
| 173Yb | CD56 | HCD56 | BioLegend | Yes | CD56 | HCD56 | BioLegend | Yes |
| 174Yb | HLA-DR | LN3 | Thermo Fisher (eBioscience) | Yes | HLA-DR | LN3 | Thermo Fisher (eBioscience) | Yes |
| 175Lu | CD54 | LB-2 | BD Biosciences | Yes | CD127 | A019D5 | BioLegend | Yes |
| 176Yb | CD117 (c-KIT) | 104D2 | BioLegend | Yes | IL32 | KU32-52 | BioLegend | No |
| 209Bi | CD11b | ICRF44 | Fluidigm | Yes | CD11b | ICRF44 | Fluidigm | Yes |

**Supplementary Table S4. Antibody panels used for CyTOF analysis.** The markers used for cell clustering by FlowSOM and ConsensusClusterPlus are provided.

### Supplementary Figure S1

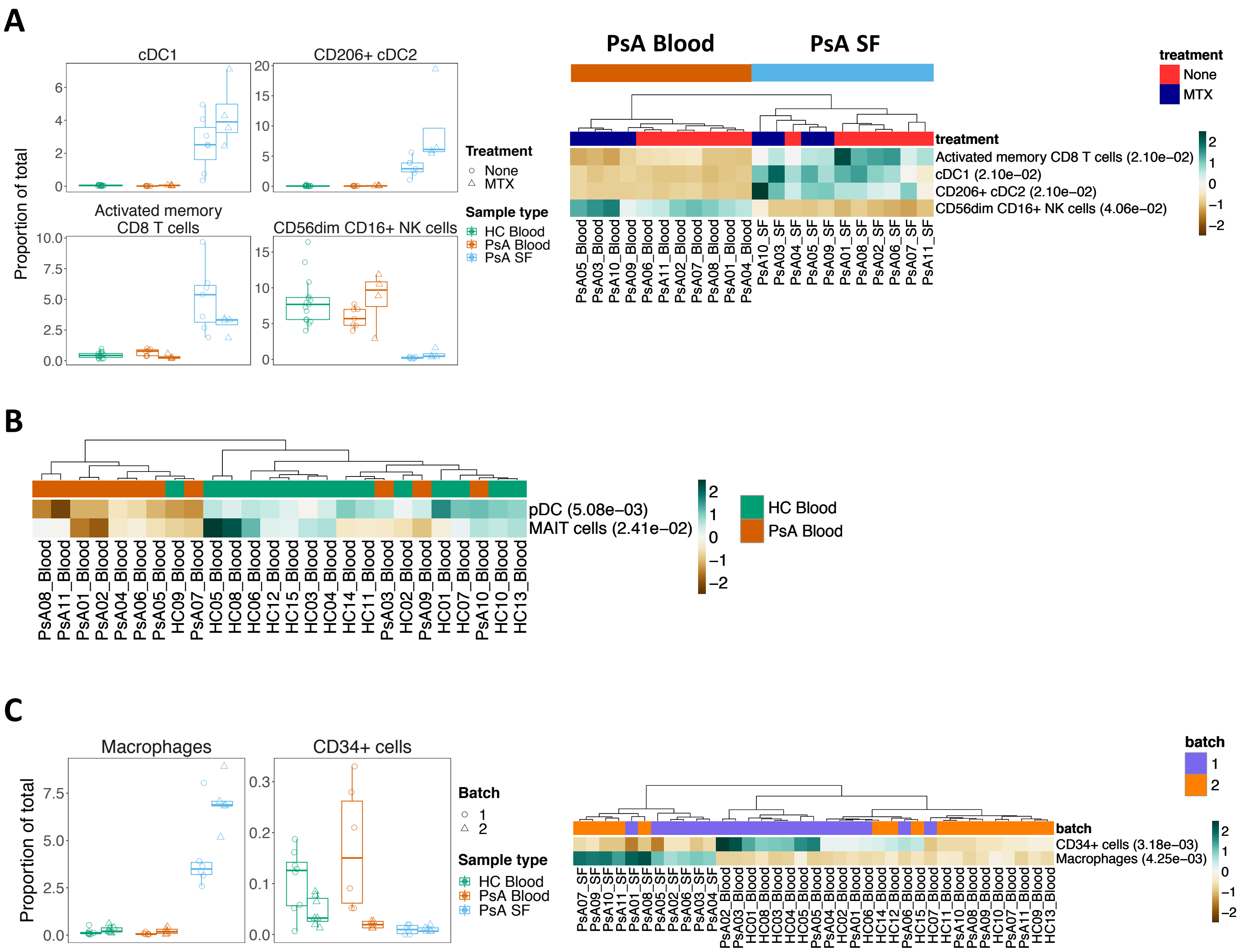


**Supplementary Figure S1. Effect of batch and treatment on cell population abundances when stained with CyTOF phenotyping panel.** (A) Comparison between PsA patients that were on no treatment or methotrexate (MTX). Differential analysis was conducted using a generalized linear mixed model (GLMM) that took into account sample type, batch, treatment, patient and pairing: y/total ~ sample type + batch + treatment + (1|sample_id) + (1|patient_id). (B) Comparison between HC blood and PsA blood groups. Differential analysis was conducted using a GLMM that took into account sample type, batch, patient and pairing: y/total ~ sample type + batch + (1|sample_id) + (1|patient_id). (C) Comparison between batch 1 and 2. Differential analysis was conducted using a GLMM that took into account sample type, batch, patient and pairing: y/total ~ sample type + batch + (1|sample_id) + (1|patient_id). The heatmaps represent arcsine-square-root transformed cell frequencies that were normalised per cluster to mean of zero and standard deviation of one. Dendrograms for samples were constructed with hierarchical clustering (Euclidean distance, average linkage). To account for the multiple testing correction, a false discovery rate cutoff of 5% was applied. Numbers in the brackets next to the cluster names indicate adjusted p-values. Any remaining neutrophils and unidentified cells were omitted from analysis.

### Supplementary Figure S2

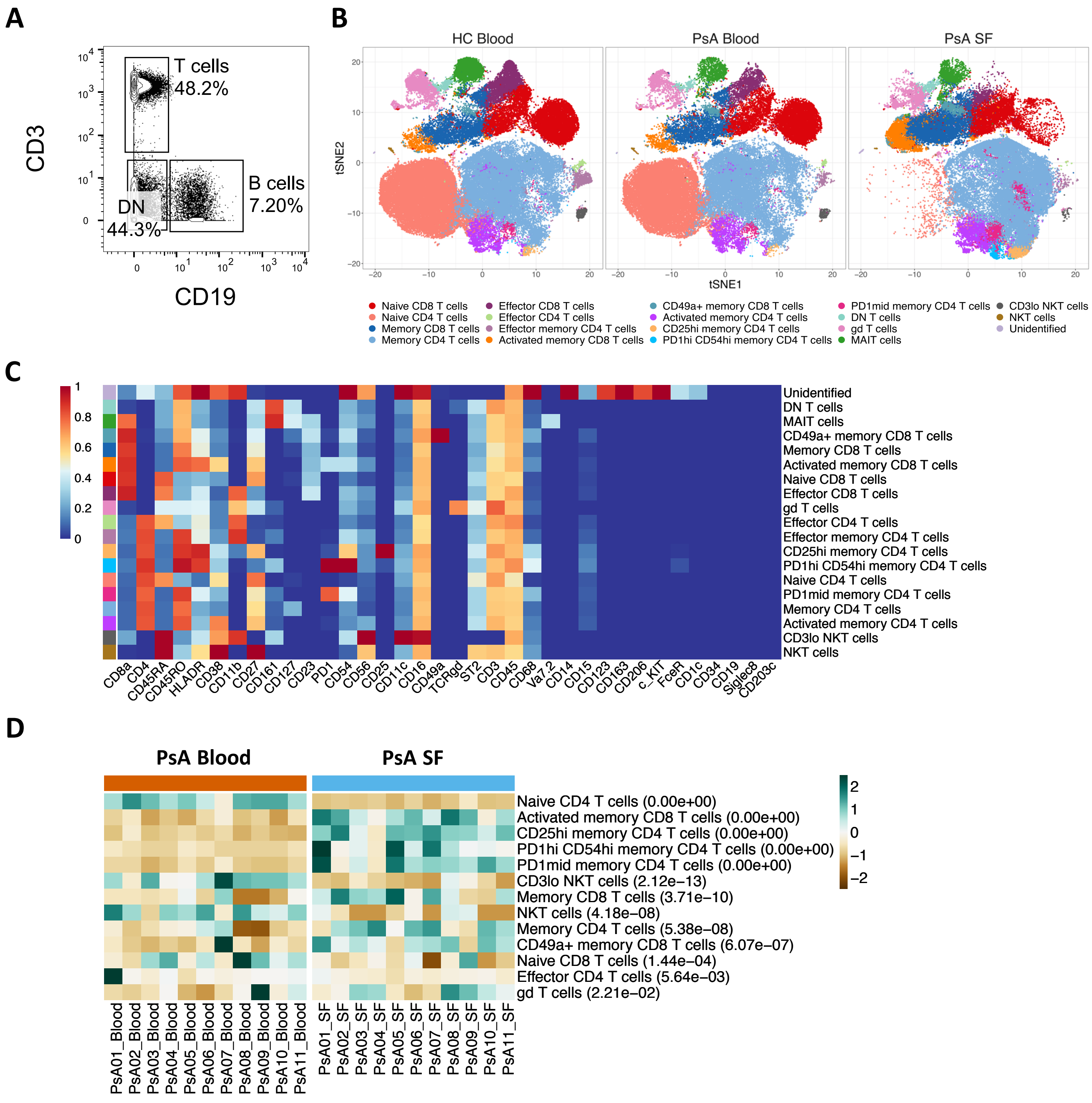


**Supplementary Figure S2. Differential analysis of CD3+ cells in blood versus SF based on a CyTOF phenotyping panel.** Samples were pre-processed to exclude all cells other than CD3+ cells using FlowJo. All samples were downsampled to an equal number of events (9,290 events). (A) Representative example of CD3+ cells used for clustering. (B) t-SNE plots based on the arcsinh-transformed expression of 36 markers in 5000 randomly selected cells from each sample. Cells are coloured according to the annotated and merged clusters. Stratified by sample type. (C) A heatmap of the median arcsinh-transformed marker intensity normalised to a 0 to 1 range across the 25 clusters obtained using FlowSOM and ConsensusClusterPlus, of the 36 markers used for clustering, following annotation and merging. (D) Comparison between PsA Blood and PsA SF groups. Differential analysis was conducted using a GLMM that took into account sample type, batch, patient and pairing: y/total ~ sample type + batch + (1|sample_id) + (1|patient_id). The heatmap represents arcsine-square-root transformed cell frequencies that were normalised per cluster to mean of zero and standard deviation of one. To account for the multiple testing correction, a false discovery rate cutoff of 5% was applied. Numbers in the brackets next to the cluster names indicate adjusted p-values.

#### Supplementary Figure S3

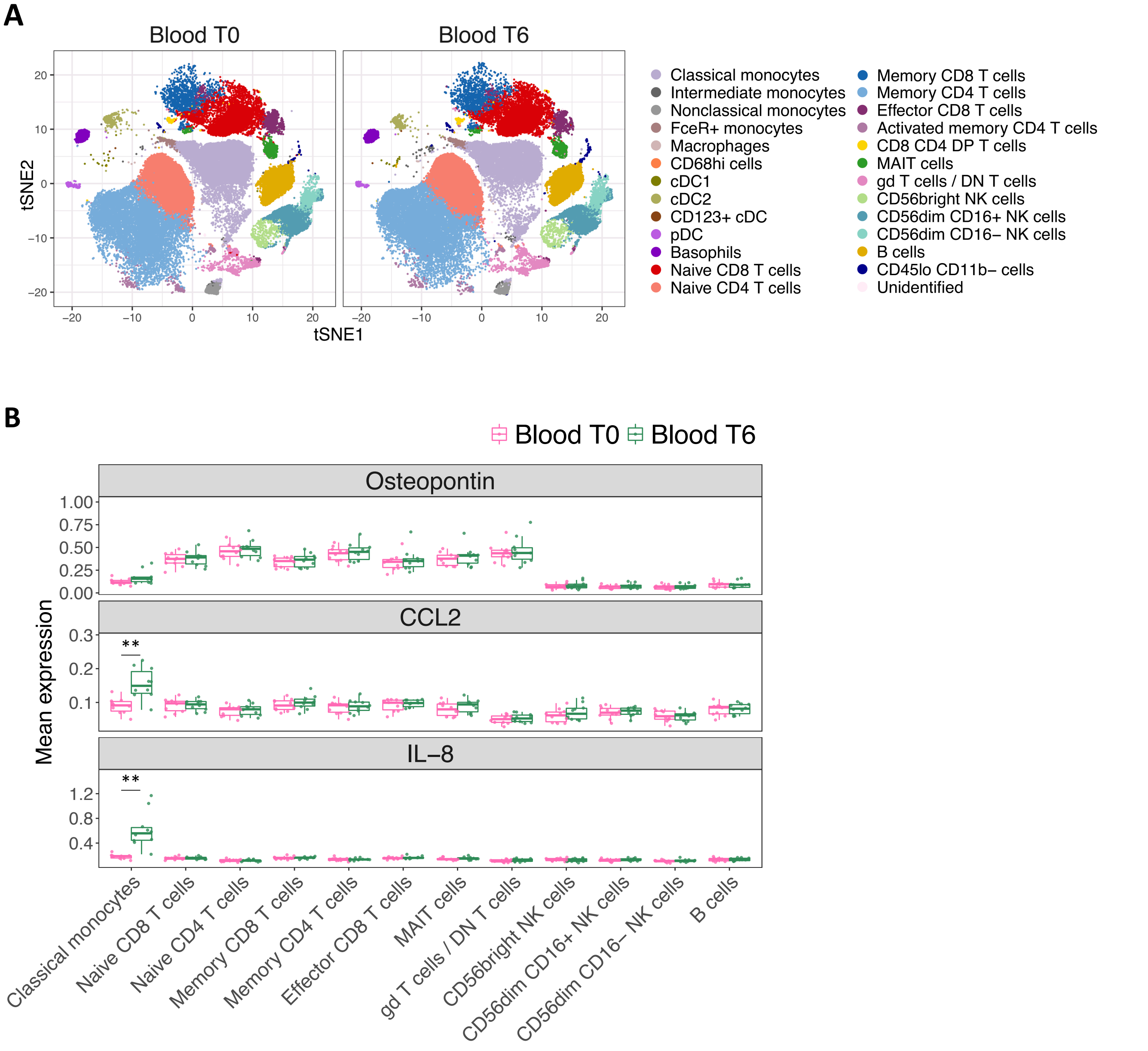


**Supplementary Figure S3. Analysis of osteopontin, CCL2 and IL-8 production by PsA blood cell populations over 6 h *ex vivo*.** (A) t-SNE plots based on the arcsinh-transformed expression of 18 markers in 5000 randomly selected cells from each sample (n = 10, only blood shown). Cells are coloured according to the annotated and merged clusters. Stratified by time. (B) Mean expression of osteopontin, CCL2 and IL-8 across the blood cell populations; any cell population containing <50 cells was omitted.

** indicates an overall increase in expression of at least 25% from T0 to T6 (with T6-T0 >0.05%), and a FDR <0.01.

### Supplementary Figure S4

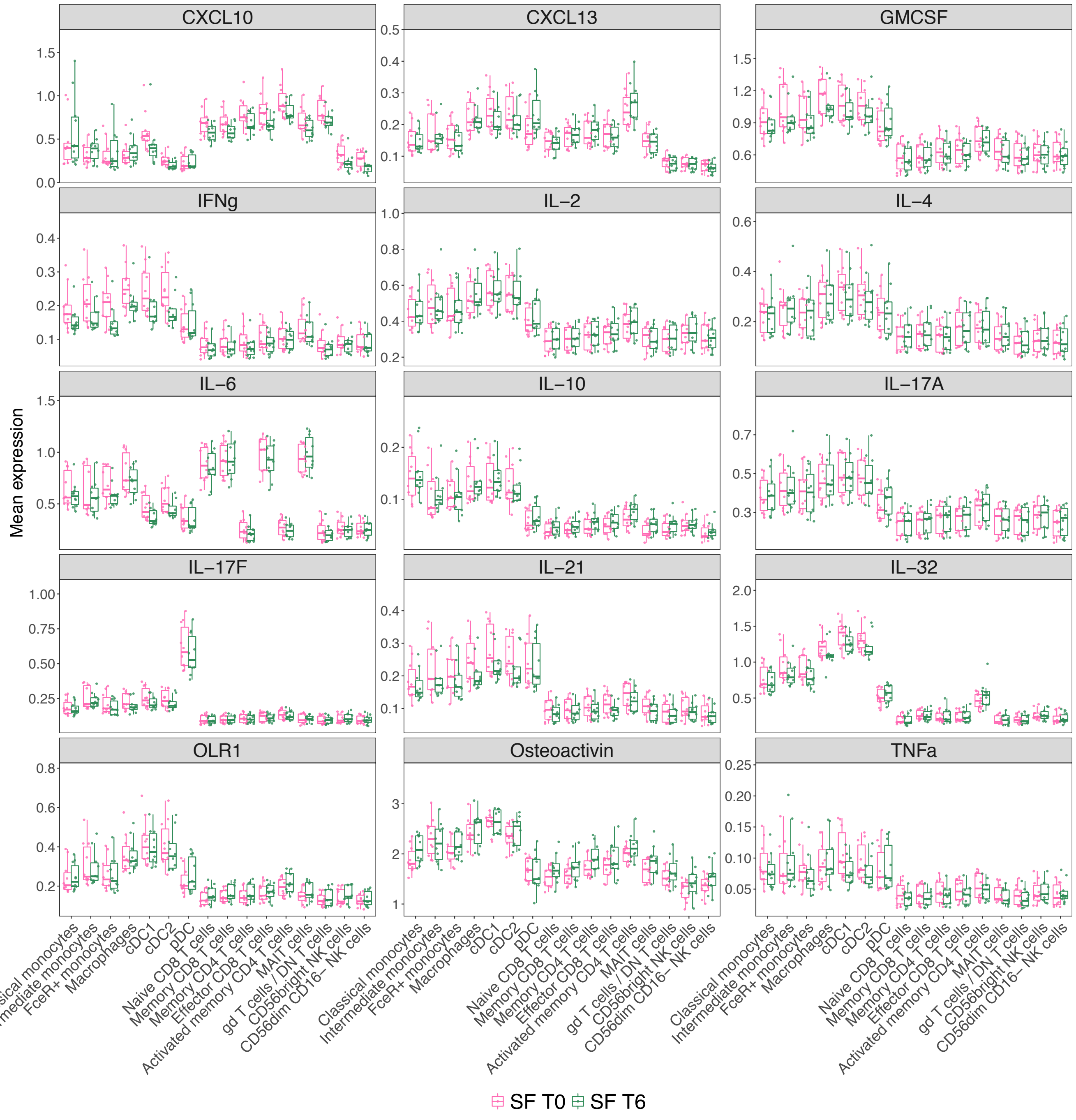


**Supplementary Figure S4. CyTOF analysis of spontaneous cytokine production by SF clusters showing mean expression of all intracellular proteins.** Any cell population containing <50 cells was omitted.
** indicates an overall increase in expression of at least 25% from T0 to T6 (with T6-T0 >0.05%), and a FDR <0.01.

### Supplementary Figure S5

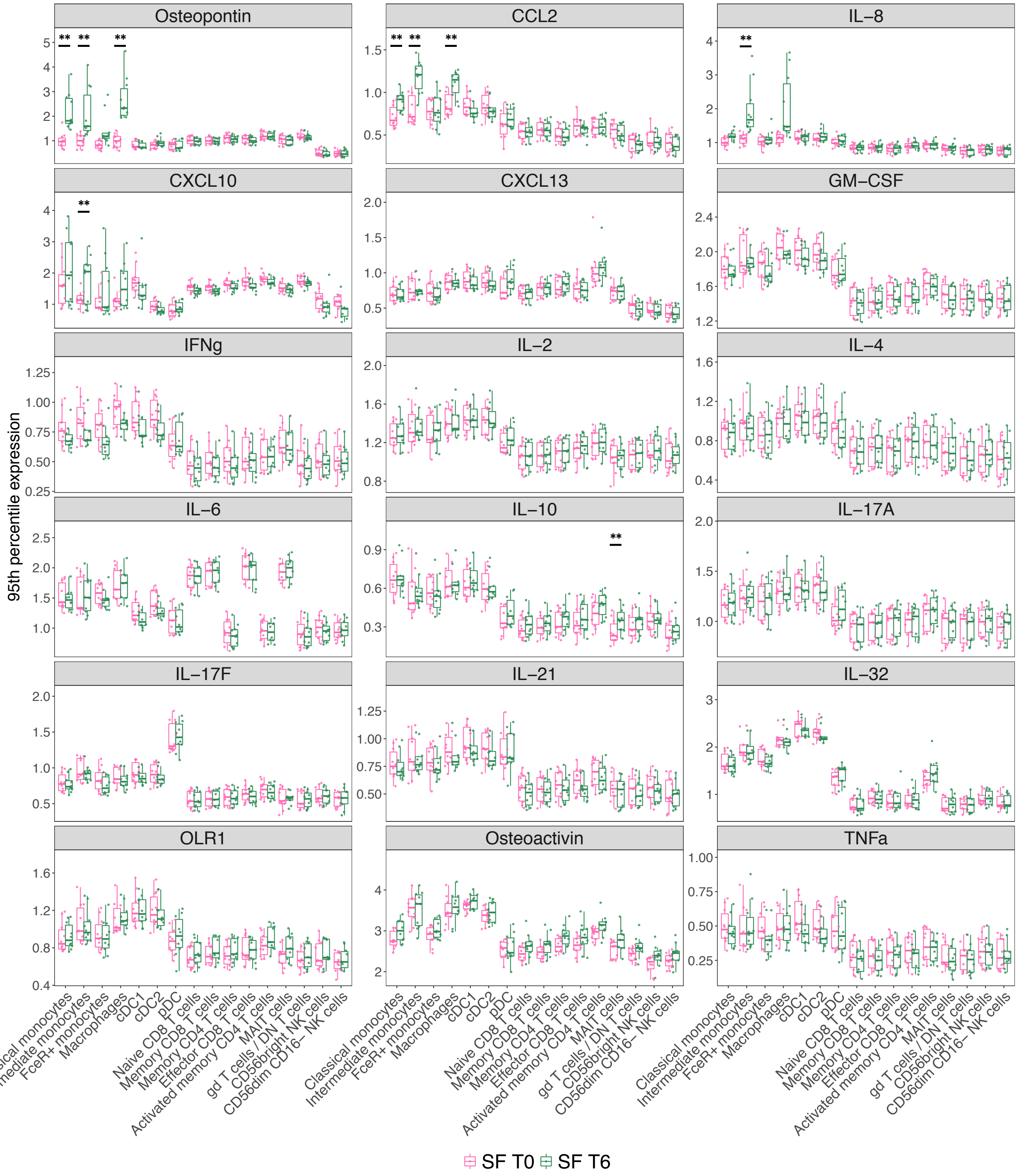


**Supplementary Figure S5. CyTOF analysis of spontaneous cytokine production by SF clusters showing 95th percentile expression of all intracellular proteins.** Any cell population containing <50 cells was omitted.
** indicates an overall increase in expression of at least 25% from T0 to T6 (with T6-T0 >0.05%), and a FDR <0.01.

### Supplementary Figure S6

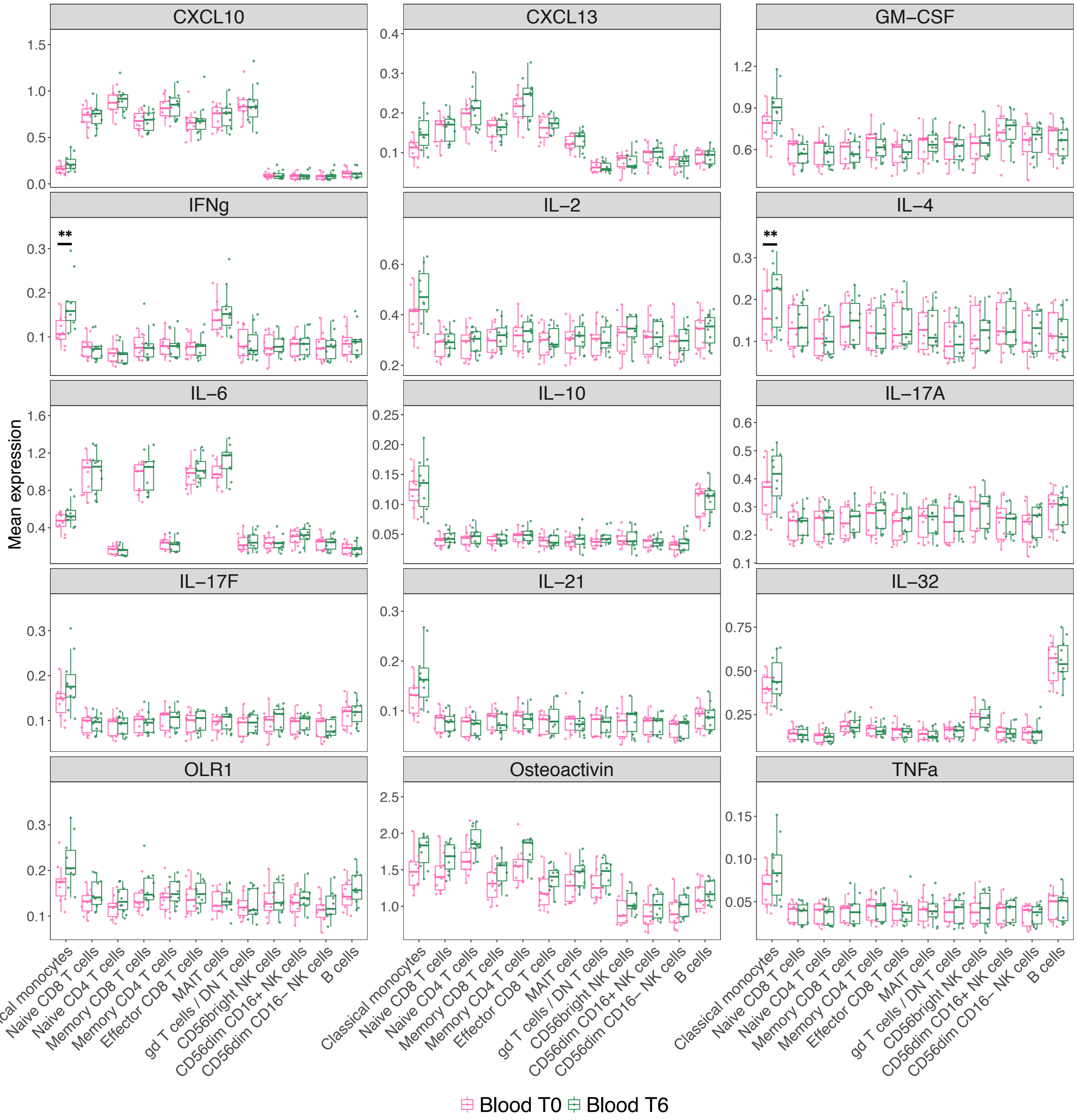


**Supplementary Figure S6. CyTOF analysis of spontaneous cytokine production by blood clusters showing mean expression of all intracellular proteins.** Any cell population containing <50 cells was omitted.

** indicates an overall increase in expression of at least 25% from T0 to T6 (with T6-T0 >0.05%), and a FDR <0.01.

### Supplementary Figure S7

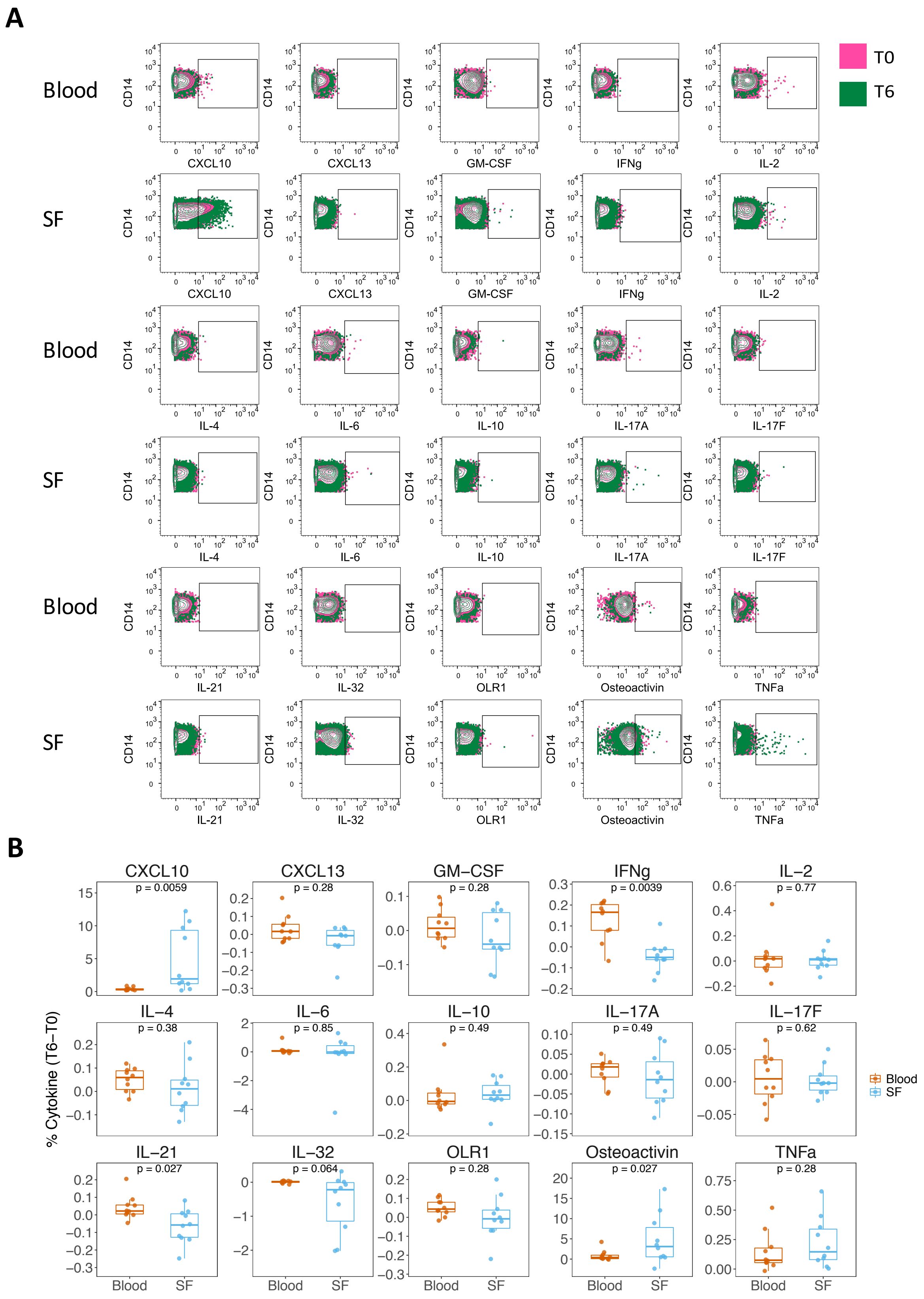


**Supplementary Figure S7. Manual CyTOF analysis of spontaneous cytokine production by blood and SF monocytes/macrophages.** (A) Following data-preprocessing, the .fcs files were gated on CD3- CD19- CD11c+ CD14+ CD123- cells using FlowJo. A representative patient is shown. (B) The percentage of intracellular protein was calculated by subtracting the amount at time T0 from time T6 per patient sample for both blood and SF. All p-values were calculated using paired Wilcox test.

### Supplementary Figure S8

CD14+ monocytes

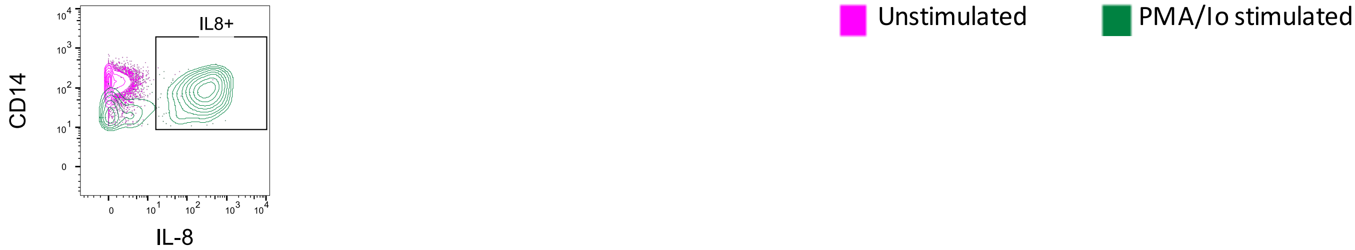


CD4+ T cells

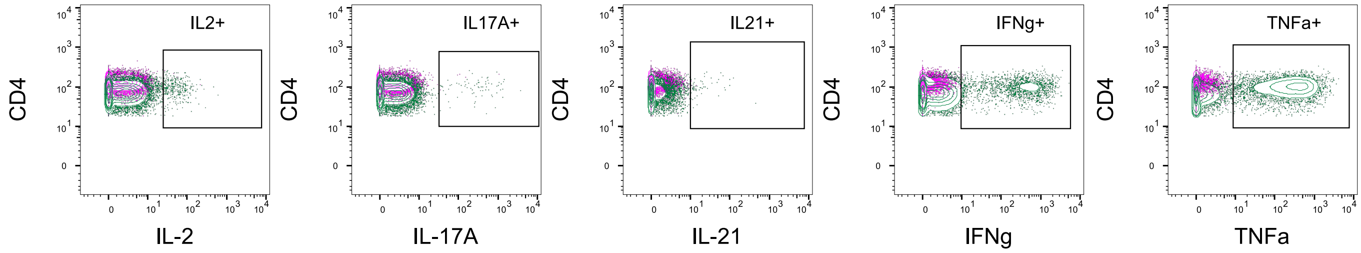


**Supplementary Figure S8. Confirmation of capacity to detect a subset of intracellular proteins by CyTOF.** Whole blood from a healthy control was stimulated for 4 h with 100 μg/ml PMA and 1 μg/ml Ionomycin in the presence of Brefeldin A/monensin. Stimulated samples were included in all 3 batches of the intracellular panel and gated in exactly the same manner as all other samples using FlowJo.

### Supplementary Figure S9

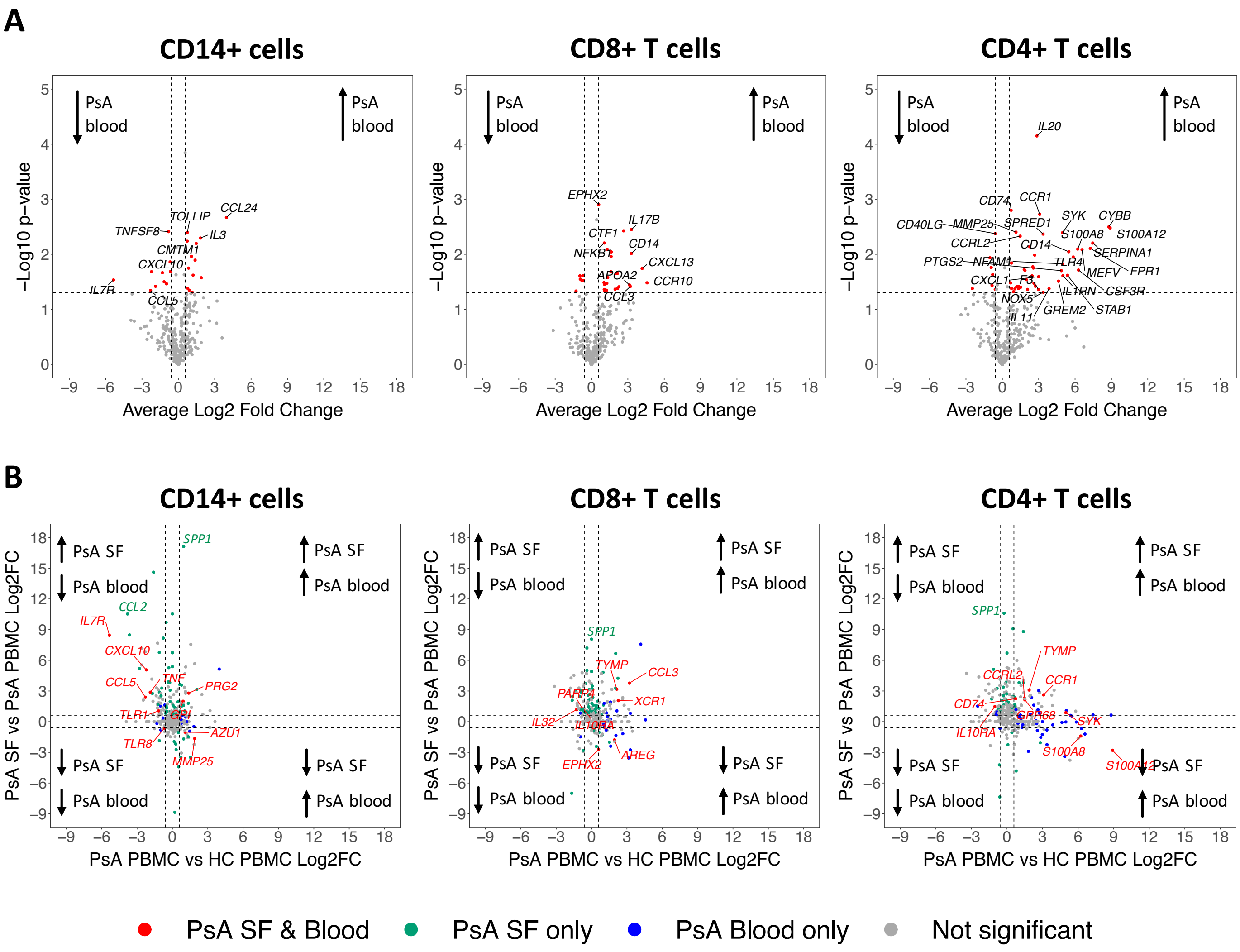


**Supplementary Figure S9. Gene expression analysis of isolated CD14+ cells, memory CD4+ T cells and memory CD8+ T cells from PsA PBMCs (n = 3) compared to healthy control (HC) PBMCs (n = 3).** (A) Volcano plots showing differences in gene expression between PsA blood and HC blood. The significance of the modulation in gene expression between PsA PBMCs compared to healthy PBMCs (y-axis) is plotted against the log2 of the mean FC (x-axis). Genes showing p-values<0.05 (one sample t-test) and mean FC>1.5 are coloured in red. Arrows indicate the direction of upregulation and downregulation of transcripts in PsA blood. (B) Comparison of gene expression across PsA tissues (SF vs blood) and against healthy controls (PsA blood vs healthy blood). Genes that were significantly up- or downregulated when comparing tissue type in PsA are in green, those significant genes that were up- or downregulated when comparing PsA blood to healthy blood are in blue, and those genes that were significant in both PsA SF compared to blood, and in PsA blood compared to healthy blood are in red.

#### Supplementary Figure S10

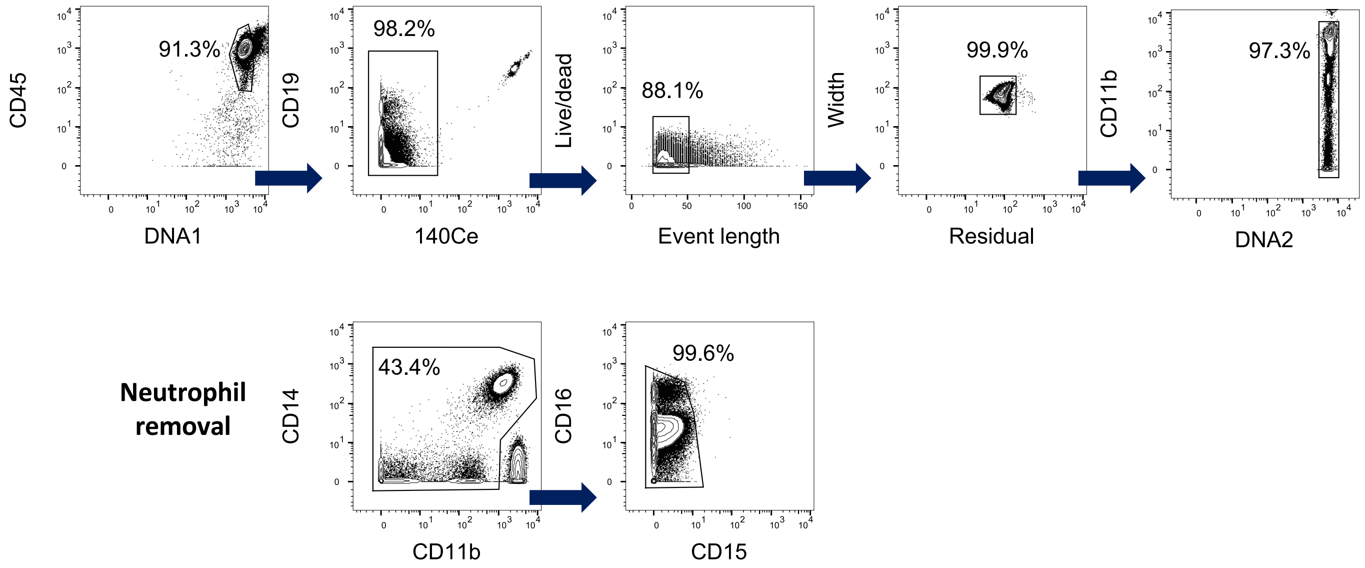


**Supplementary Figure S10. Gating strategy for pre-processing CyTOF files.** Data were pre-processed to remove debris, normalization beads, doublets and non-specific staining. Neutrophils were removed at this stage as they represented the majority of cells and so may bias the clustering. Biaxial manual gating was performed using FlowJo software (representative phenotyping panel-stained sample shown).

### Supplementary Figure S11

**A**

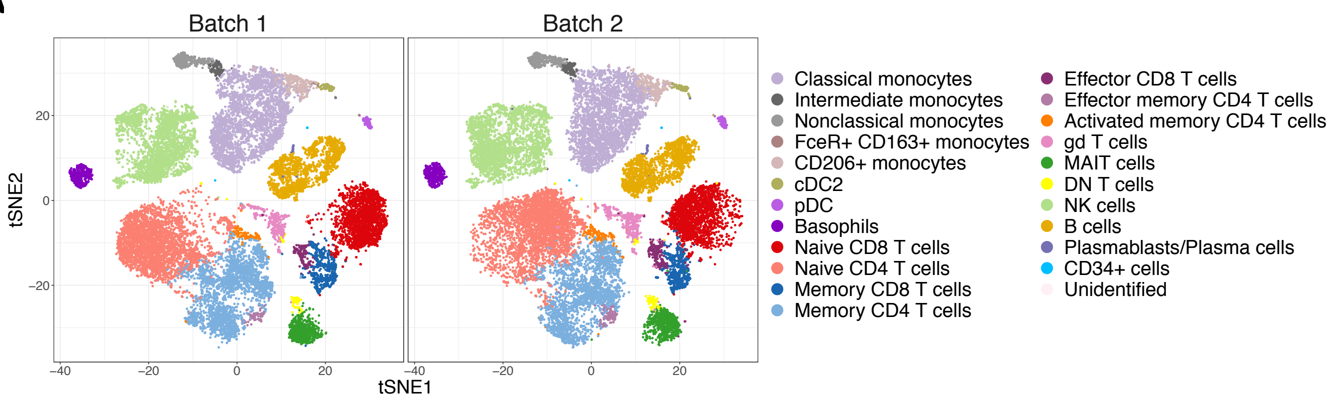


**B**

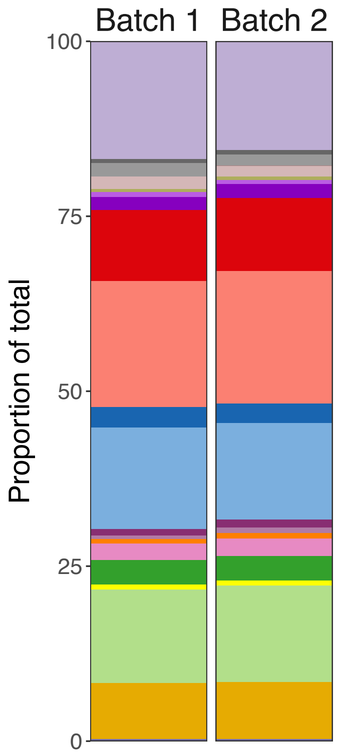


| Cluster | Batch 1 proportion | Batch 2 proportion |
|---|---|---|
| Classical monocytes | 16.80 | 15.59 |
| Intermediate monocytes | 0.60 | 0.60 |
| Nonclassical monocytes | 1.90 | 1.59 |
| FceR+ CD163+ monocytes | 0.03 | 0.05 |
| CD206+ monocytes | 1.75 | 1.53 |
| cDC2 | 0.48 | 0.45 |
| pDC | 0.68 | 0.60 |
| Basophils | 1.90 | 1.99 |
| Naive CD8 T cells | 10.15 | 10.38 |
| Naive CD4 T cells | 17.95 | 18.93 |
| Memory CD8 T cells | 2.96 | 2.86 |
| Memory CD4 T cells | 14.47 | 13.74 |
| Effector CD8 T cells | 0.96 | 1.18 |
| Effector memory CD4 T cells | 0.47 | 0.77 |
| Activated memory CD4 T cells | 0.66 | 0.79 |
| gd T cells | 2.38 | 2.54 |
| MAIT cells | 3.51 | 3.46 |
| DN T cells | 0.69 | 0.68 |
| NK cells | 13.39 | 13.84 |
| B cells | 7.97 | 8.14 |
| Plasmablasts/Plasma cells | 0.21 | 0.21 |
| CD34+ cells | 0.08 | 0.02 |
| Unidentified | 0.01 | 0.04 |

**Supplementary Figure S11. Comparison of CyTOF data from the 2 phenotyping panel batches using a technical control**. To observe any batch effects, HC06 was included in both batches of the phenotyping panel. (A) t-SNE plots based on the arcsinh-transformed expression of 36 markers in 20,000 randomly selected cells from each batch. Cells are coloured according to the annotated and merged clusters. (B) Cell composition frequencies stratified by batch; neutrophils that were still present were omitted from this analysis.

### Supplementary Figure S12

**A**

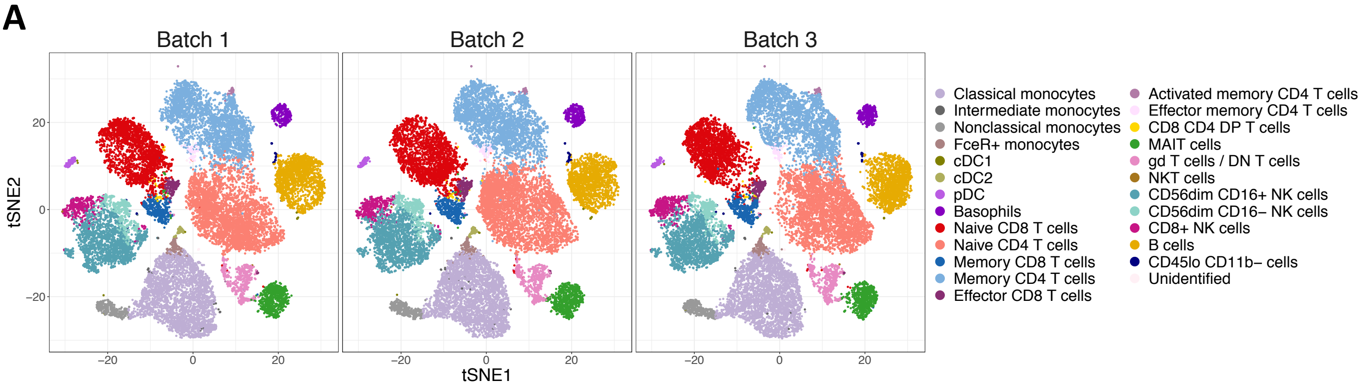


**B**

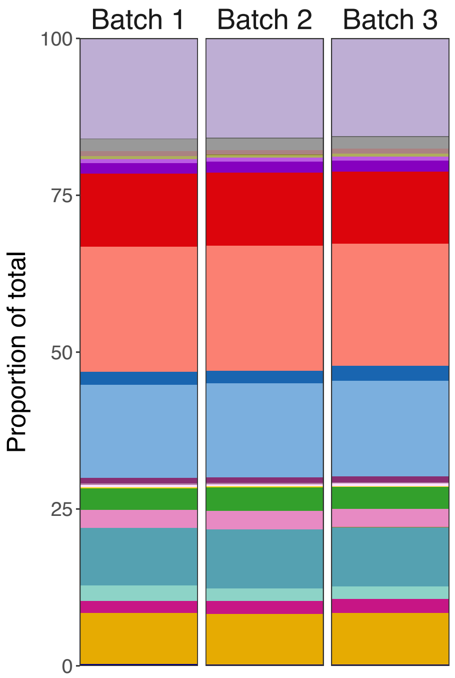


| Cluster | Batch 1 proportion | Batch 2 proportion | Batch 3 proportion |
|---|---|---|---|
| Classical monocytes | 15.94 | 15.86 | 15.62 |
| Intermediate monocytes | 0.14 | 0.16 | 0.16 |
| Nonclassical monocytes | 1.89 | 1.80 | 1.79 |
| FceR+ monocytes | 0.79 | 0.75 | 0.78 |
| cDC1 | 0.036 | 0.035 | 0.043 |
| cDC2 | 0.50 | 0.46 | 0.44 |
| pDC | 0.60 | 0.58 | 0.61 |
| Basophils | 1.69 | 1.76 | 1.81 |
| Naive CD8 T cells | 11.65 | 11.69 | 11.41 |
| Naive CD4 T cells | 19.90 | 19.90 | 19.53 |
| Memory CD8 T cells | 2.06 | 2.05 | 2.35 |
| Memory CD4 T cells | 14.87 | 14.93 | 15.28 |
| Effector CD8 T cells | 0.89 | 0.95 | 0.89 |
| Activated memory CD4 T cells | 0.20 | 0.19 | 0.22 |
| Effector memory CD4 T cells | 0.41 | 0.36 | 0.41 |
| CD8 CD4 DP T cells | 0.16 | 0.15 | 0.13 |
| MAIT cells | 3.41 | 3.70 | 3.55 |
| gd T cells / DN T cells | 2.90 | 3.00 | 2.87 |
| NKT cells | 0.016 | 0.018 | 0.022 |
| CD56dim CD16+ NK cells | 9.13 | 9.35 | 9.37 |
| CD56dim CD16- NK cells | 2.52 | 2.05 | 2.07 |
| CD8+ NK cells | 1.85 | 2.07 | 2.22 |
| B cells | 8.18 | 8.05 | 8.19 |
| CD45lo CD11b- cells | 0.21 | 0.18 | 0.19 |
| Unidentified | 0.054 | 0.005 | 0.021 |

**Supplementary Figure S12. Comparison of CyTOF data from the 3 intracellular panel batches using a technical control.** To observe any batch effects, HC06 was included in all 3 batches of the intracellular panel. (A) t-SNE plots based on the arcsinh-transformed expression of 18 lineage markers in 20,000 randomly selected cells from each batch. Cells are coloured according to the annotated and merged clusters. (B) Cell composition frequencies stratified by batch.

### Supplementary Figure S13

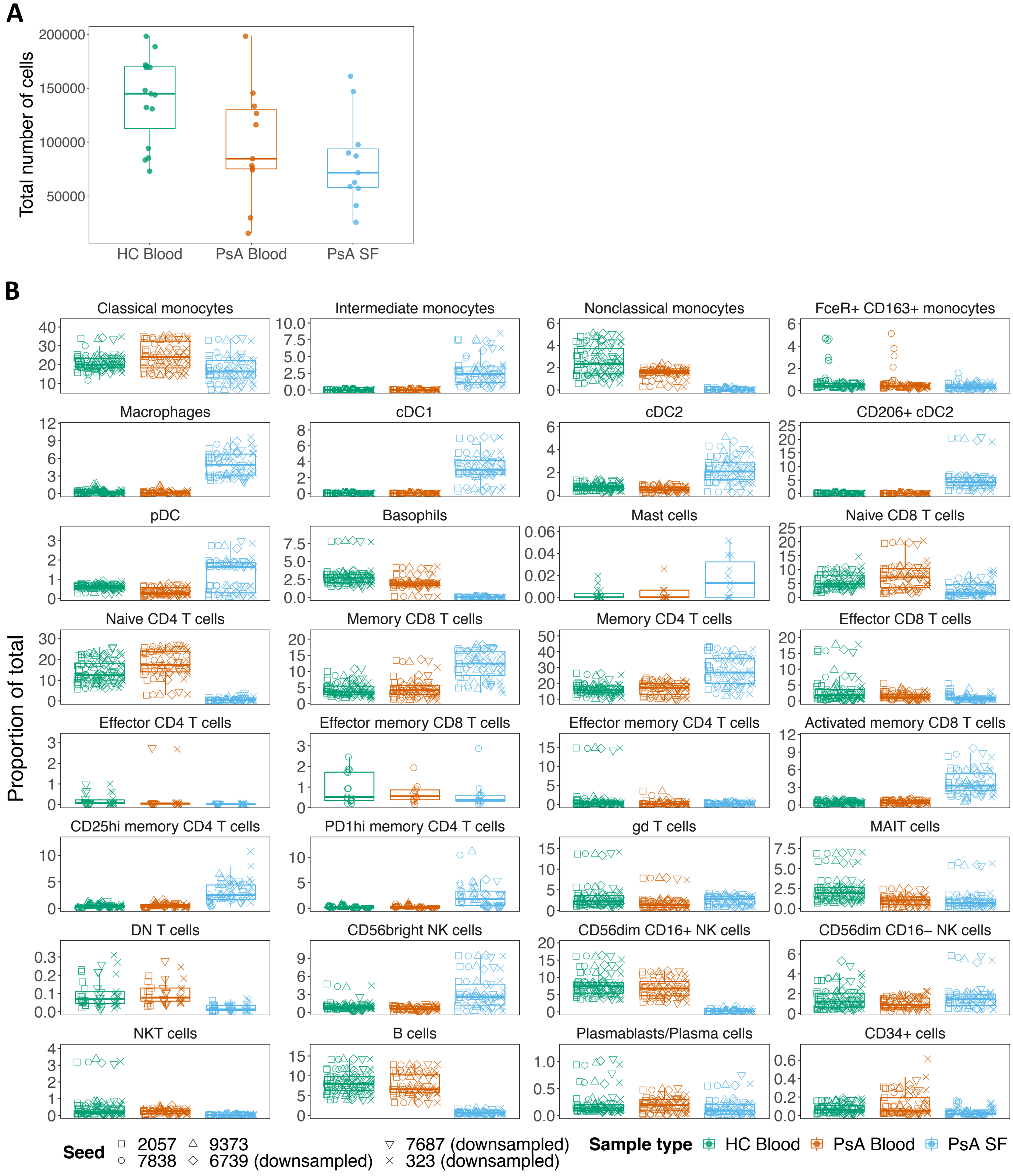


**Supplementary Figure S13. Effect of seed and total number of cells on cell population abundances when stained with CyTOF phenotyping panel.** (A) Total number of cells per sample following pre-processing but prior to downsampling. (B) Three random seeds from full pre-processed files and three seeds from downsampled files (downsampled to the sample with the fewest events) were run to determine whether multiple random starts affected cell population abundance. While some rarer cell populations were identified with some seeds, we continued with seed 6739 for our full analysis as this consistently returned similar abundances to the other seeds tested.
